# *Visual circuitry for distance estimation in* Drosophila

**DOI:** 10.1101/2024.12.25.630346

**Authors:** Joseph Shomar, Elizabeth Wu, Braedyn Au, Kate Maier, Baohua Zhou, Natalia C.B. Matos, Garrett Sager, Gustavo Santana, Ryosuke Tanaka, Caitlin Gish, Damon A. Clark

## Abstract

Animals must infer the three-dimensional structure of their environment from two-dimensional images on their retinas. In particular, visual cues like motion parallax and binocular disparity can be used to judge distances to objects. Studies across several animal models have found neural signals that correlate with visual distance, but the causal role of these neurons in distance estimation as well as the range of possible neural properties that can inform distance estimation have remained poorly understood. Here, we developed a novel high-throughput behavioral assay to identify neurons in the *Drosophila* visual system that are involved in distance estimation during free locomotion. We found that silencing the primary motion detectors in the fly visual system eliminated their ability to perceive distance, consistent with a reliance on motion parallax to judge distance. Through a targeted silencing screen of visual neurons during behavior and through *in vivo* two-photon microscopy, we identified a visual projection neuron that encodes the parallax signal in the relative motion of foreground and background. Interestingly, it differs from previously identified parallax-tuned neurons in its lack of direction selectivity both to moving bars and to moving backgrounds. This non-canonical tuning is interpretable in the context of parallax signals that the fly would likely encounter during naturalistic walking behavior. Our results demonstrate how both direction selective and non-direction selective feature-detecting neurons can contribute to distance estimation using parallax cues, providing a framework for considering broader classes of parallax-encoding neurons in distance estimation across visual systems.

## Introduction

As animals navigate, it is often critical to determine the three-dimensional structure of the world. Using vision, the problem is to infer three-dimensional structure from two-dimensional projections of the world onto the retina, a process known as spatial vision^1^. The inference problem is non-trivial because many three-dimensional structures correspond to a single retinal image (**Fig. 1A**), but the problem is solved by the brains of many animals across phyla^2–8^. To solve the problem of spatial vision, animals use a variety of visual cues^9^, including relative size^10,11^, occlusion^11,12^, binocular vergence^13^, optic flow^14–16^, binocular disparity^17–21^, and motion parallax^17,21–26^.

**Figure 1.**
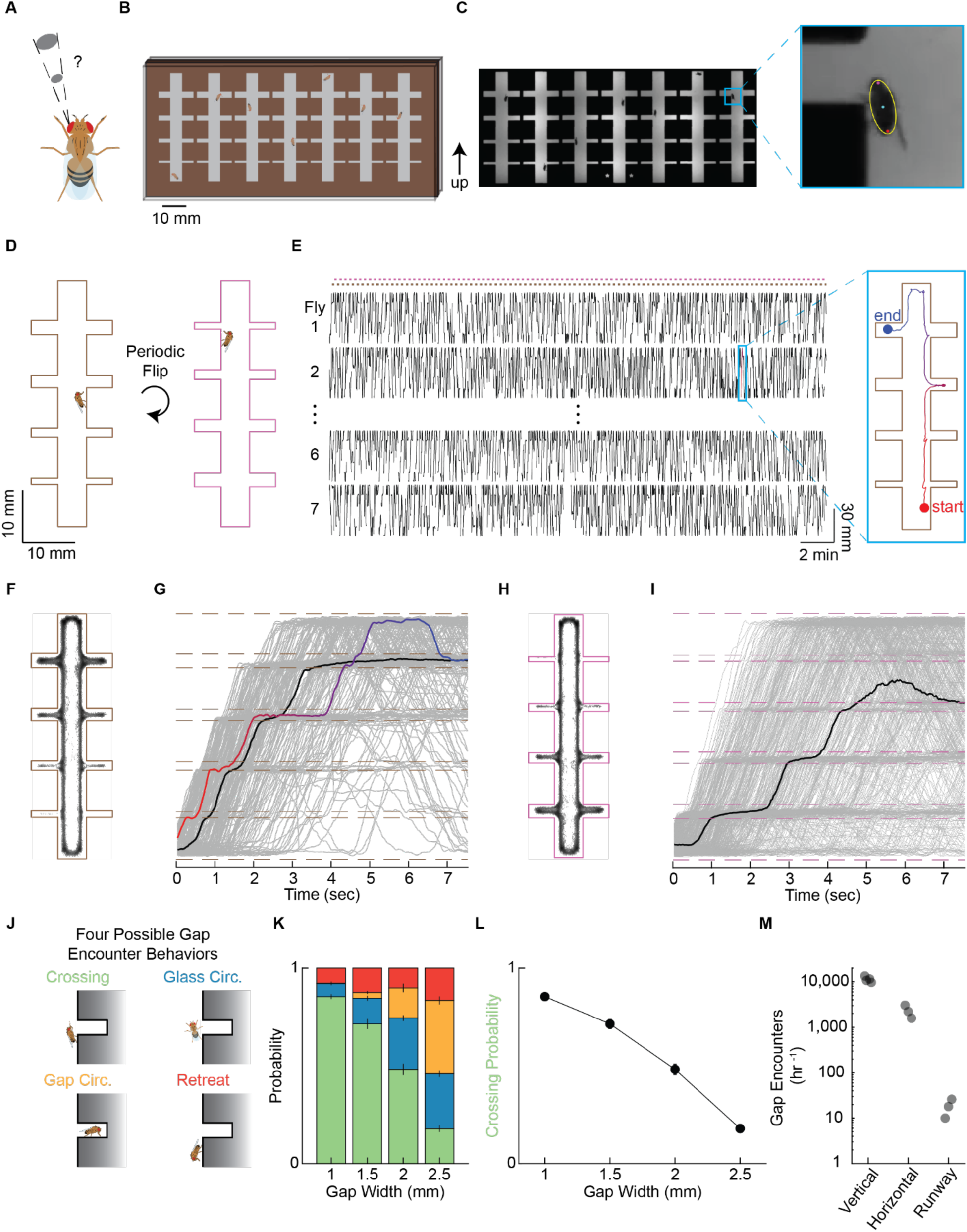
High-throughput gap crossing assay. A) Spatial vision is inherently ambiguous, since two-dimensional images on the retina can reflect an infinite number of three-dimensional configurations of the world. B) Diagram of high-throughput gap crossing assay design. Male flies were individually placed in laser cut corridors and sandwiched between coated glass panes. C) Example image from a gap crossing experiment. Inset: Overlaid tracked features for a single fly (head in magenta, centroid in cyan, rear in red, body as fitted ellipse in yellow). D) Diagrams of corridor in two flipped orientations. E) Vertical position of wild type (WT) flies over the course of a single experiment. Orientation of corridor in brown (orientation 1) and pink (orientation 2). Inset: Two-dimensional trajectory of a single fly within a corridor during a single trial. F) Location of all WT fly centroids within a corridor in orientation 1. N = 28 flies here and in panels (G), (H), (I), (K), and (L). G) Vertical position trajectories within each trial for WT flies in orientation 1, selected for those starting below the first gap, showing a random sample (gray), median trajectory (black), and the trajectory shown in (E) (red to blue). Gap locations are denoted by dashed lines. H) As in (F) for orientation 2. I) As in (G) for orientation 2. J) Diagrams of the four possible gap encounter behaviors that a fly can perform upon reaching a gap (crossing, glass circumvention, gap circumvention, and retreat). K) Probability of each gap encounter behavior among WT flies for each gap size. L) Probability of crossing among WT flies for each gap size. M) Number of gap encounters per hour performed by WT flies in three different gap crossing assays. The runway assay is equivalent to that used in previous studies^25^. All error bars represent one standard error of the mean.

The cues of binocular vergence, binocular disparity, and motion parallax signals are particularly important for visual distance estimation across many animals^25,27–30^. Binocular vergence corresponds to the degree of inward rotation of a binocular observer’s eyes when fixating on an object^31^, whereas binocular disparity arises from the different images formed on the retinas of two spatially separated eyes^32^. In comparison, motion parallax is a geometric phenomenon that occurs when an observer translates through the world — in that case, the relative speed of objects on the observer’s retina depends on the relative distance of the objects, with more distance objects moving slower than closer ones^32^. Under certain conditions, binocular disparity and motion parallax are computationally analogous^32^. Since many animals successfully determine the structure of their surrounding world using these cues, identifying behaviorally relevant neurons and determining how they encode these distance cues is crucial for understanding how animals solve the problem of spatial vision.

The neural encoding of visual distance has been studied in a variety of models, including non-human primates^17,21,23,24,33^, mice^34,35^, gerbils^6^, birds^11,36^, locusts^22,37^, and mantids^12,19,20,38,39^. In primates, cortical area MT contains many direction- and disparity-selective neurons thought to be involved in distance estimation tasks^17,23,24,33,40^. Reversible inactivation of area MT eliminates a trained monkey’s ability to perform some, but not all, distance-based tasks^33^. Similarly, optogenetic suppression of the primary visual cortex in mice degrades performance on distance estimation tasks^35^. Locusts make use of lateral peering movements to amplify motion parallax signals^37^, and manipulating motion parallax signals leads to predictable changes in their distance-dependent behaviors^22,37^. Praying mantids make use of both monocular and binocular cues to estimate distances^12,18–20,38,39^, and new anaglyph techniques and electrophysiological recordings have identified disparity-selective neurons in the mantid brain^18,20,39^. Collectively, these studies have identified a host of neurons that encode distance cues as well as brain regions required for distance estimation behaviors. Still, causal contributions of specific cell types that encode visual distance cues to distance-dependent behaviors are yet to be established^32^.

The genetic toolkit in *Drosophila* makes it possible to causally connect individual classes of neurons to behavior^41^. Flies exhibit various distance-dependent behaviors and neural signals^14,15,25,26,42–44^ and have been proposed to exploit cues such as binocular disparity, motion parallax, and optic flow^14,15,25,26,44^. Gap crossing is a powerful paradigm for observing visual distance estimation in flies^25,42–44^ and in other animal models^6,35^, and can be used to test the relative importance of different visual cues^6,25,44^. In the gap crossing paradigm, an animal is presented with a gap in the surface of its walking path. The frequency of crossing depends on the gap size and, critically, on visual inputs. In *Drosophila*, gap crossing experiments combined with genetic silencing have identified two neurons that alter distance-dependent behavior^42^, although it remains unclear how they encode information relevant for distance estimation. Historically, gap crossing assays have been limited by either very low-throughput^25^ or coarse measurements of behavior in high-throughput experiments^42^. Here, we have developed a novel behavioral assay to collect high-throughput and finely tracked gap crossing behavior in *Drosophila*.

Using this high-throughput behavioral assay, we have characterized gap crossing behavior in flies, first investigating the relative use and source of specific visual cues. By conducting a targeted genetic silencing screen of many feature-selective neurons across the optic lobe, we identified neuron classes required for wild type gap crossing behavior. These experiments showed for the first time that the primary local motion detectors in the fly visual system, T4 and T5, are required for visual distance estimation. A second feature-encoding neuron identified in the screen was LC15, a visual projection neuron, which we subsequently characterized using *in vivo* two-photon calcium imaging. We discovered that though it lacks canonical features for motion parallax selectivity, it nonetheless encodes the relative distance between objects and backgrounds based on motion parallax cues. Overall, these results identify circuitry that performs parallax-based distance estimation in flies and reveals a breadth in potential response properties that can encode relative object distances.

## Results

### A high-throughput gap crossing assay

To use a gap crossing paradigm to investigate the neural basis of distance estimation, we first designed a new, high-throughput gap crossing assay. In this assay, individual flies are placed into separate, laser-cut corridors that contain four gaps of different sizes (1, 1.5, 2, 2.5 mm). Multiple vertically oriented corridors were enclosed in a cassette sandwiched between two transparent, glass panes (**Fig. 1B**). The panes were coated with a bitterant-infused wax to deter flies from walking on the glass (**Methods**). Infrared light illuminated flies for tracking over the course of each experiment (**Fig. 1C**), while green LEDs illuminated the rig so that the flies could see the corridors and gaps. To permit use of thermogenetic tools, the rig was heated, which also increased fly locomotion. To further increase gap encounters by flies, we took advantage of fly gravitaxis^45^, in which flies tended to walk up walls against the force of gravity. The cassette was flipped 180° (**Fig. 1D**) every eight seconds by a servo-motor to increase the rate of gap encounters. Fly position and orientation were tracked in each corridor for the entirety of the experiment (**Fig. 1E, F**). Flies that started at the bottom of a corridor tended to navigate to the top of the corridor before the next flip eight seconds later (**Fig. 1H, J**). Flies rarely entered small gaps and frequently entered larger gaps (**Fig. 1G, I**).

Upon encountering a gap, flies exhibited four distinct behaviors: gap crossing, glass circumvention, gap circumvention, or retreats (**Fig. 1J**). During a gap crossing, a fly traversed across a gap from one side to the other. During a glass or gap circumvention, a fly navigated to the other side of the gap by walking on the glass or walking into the gap, respectively. During a retreat, a fly reached a gap, then turned around and walked away from the gap. Gap encounters were initially identified by tracking the location of each fly relative to the gaps in the corridor (**Fig. S1A**). A custom image processing pipeline categorized behaviors into the four distinct categories (**Fig. S1B–G, Methods**).

The probabilities of each behavioral category depended on the size of the gap (**Fig. 1K, L**), with flies crossing smaller gaps more frequently than larger ones, consistent with prior behavioral measurements^25^. Gap crossing frequency mirrors attempt frequency and similarly depends on gap size^25^, so that crossing frequency can assay the ability to estimate the size of the gaps they encounter^42^. Since the frequency of gap crossing has a larger dynamic range than attempt frequency, behavioral data is often summarized by gap crossing frequency at each gap size (**Fig. 1L**)^25,42^.

Our rig collected ∼1,000 times as many gap encounters per hour as classical runway assays that measure crossings in one fly at a time (**Fig. 1M**). The vertical orientation of the corridors and gaps yielded approximately twice the encounters as a horizontal orientation (**Fig. 1M**), while the crossing frequencies were comparable (**Fig. S1H**). The increases in the data volume per hour yielded strong statistical power in our assays and enabled a targeted screen of neurons.

### Local motion detectors are required for gap crossing but binocular vision is not

First, to verify that the distance-dependent crossing behaviors were visually mediated, we turned off the green light. Under these conditions, the gap crossing frequency curve shifted significantly toward the smaller gaps (**Fig. 2A**), consistent with prior findings^25^. The gap crossing frequency in the dark still depends on gap size, again consistent with prior work^25^. This is likely because the flies in the dark incidentally step over the smaller but not the larger gaps.

**Figure 2.**
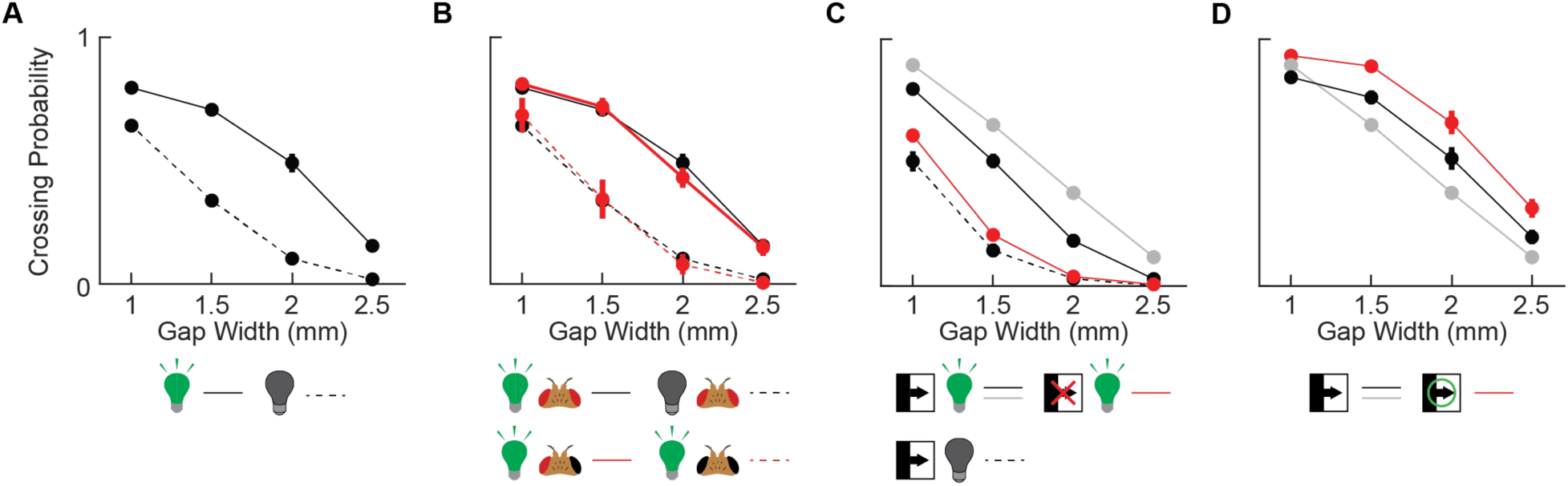
Local motion detector neurons T4 and T5 are required for gap crossing but binocular vision is not. A) Mean crossing probability for each gap size among WT flies with light (solid) and in the dark (dashed). B) Mean crossing probability for each gap size among WT flies with light (solid black), in the dark (dashed black), with light and with one eye painted (solid red), and with light and with both eyes painted (dashed red). C) Mean crossing probability for each gap size for flies with T4 and T5 silenced (red, T4T5 > shi), both parental controls (gray, T4T5 > + and solid black, empty Gal4 > shi), and one parental control in the dark (dashed black, empty Gal4 > shi). D) Mean crossing probability for each gap size for flies with T4 and T5 activated (red, T4T5 > dTRPA1) and for both parental controls (gray, T4T5 > + and solid black, empty Gal4 > dTRPA1). N = 19 to 28 flies for each genotype. All error bars represent one standard error of the mean.

Since several studies have suggested that binocular cues may be important for insect distance estimation^18–20,38,39,44^, we wanted to test whether binocular vision was necessary in our paradigm. To eliminate binocular cues, we painted over one eye of wild type flies using black nail polish and ran them in our assay. The gap crossing frequency curve of the flies with a single eye painted was statistically indistinguishable from the wild type curve (**Fig. 2B**). To verify that the nail polish had successfully prevented the flies from seeing out of the painted eye, we also painted both eyes using the same black nail polish. In that case, the gap crossing frequency curve was statistically indistinguishable from the curve of wild type flies in the dark (**Fig. 2B**), indicating that the painting had successfully blinded the flies. Together, these experiments demonstrate that binocular cues are dispensable for distance estimation in this paradigm, consistent with prior findings^25^.

Many accounts of monocular distance estimation in flies and other insects implicate the use of motion parallax^14,22,25,26,37,42^. We reasoned that if flies were using motion parallax to judge distance, then the primary motion detectors, T4 and T5^46^, were likely to be involved. To test this hypothesis, we silenced T4 and T5 neurons using shibire^ts^, which conditionally blocks synaptic transmission^47^. When T4 and T5 were silenced, the gap crossing frequency curve shifted towards smaller gap sizes (**Fig. 2C**), demonstrating that T4 and T5 are required for distance estimation during gap crossing. The geometry of motion parallax dictates that nearby objects move faster as an observer translates. Thus, silencing T4 and T5 would make it seem as if all objects had no motion across the fly’s retina and were thus far away, leading to an overestimate of gap size and reducing the crossing frequency of large gaps.

Using the same reasoning, we hypothesized that higher baseline activity in T4 and T5 could make objects appear to move faster, and hence appear closer to the fly. To test this hypothesis, we expressed in T4 and T5 the channel dTRPA1, which constitutively activates neurons at high temperatures^48,49^. When this channel was expressed, the gap crossing frequency curve shifted to larger gap sizes (**Fig. 2D**), the opposite direction from silencing the cells. This is consistent with the hypothesis that higher activity in T4 and T5 makes gaps appear smaller. In this case, flies appear capable of crossing larger gaps but modulate their attempt frequency based on perceived depth, consistent with prior suggestions^42^. Combined with the T4 and T5 silencing experiment, this data indicates that T4 and T5 can help generate motion parallax signals for distance estimation.

### Distinct behaviors yield different motion parallax signals during gap crossing

Since motion parallax is central to gap crossing behaviors, we set out to examine which aspects of fly locomotion result in parallax cues. To collect the third dimension of location information unavailable in our high-throughput assay, we designed a low-throughput assay to measure the fly’s lateral position relative to the gap. The assay had a corridor with a single gap on one side and a 45° mirror on the other (**Fig. 3A**). We identified three likely sources of motion parallax cues during locomotion towards a gap: lateral movements of the head and body^22^, bobbing movements of the head as a result of stepping^50^, and motion of the head into the gap^35^ (**Fig. 3B–D**). We used markerless tracking^51^ to help identify the head and rear of the fly in this setup (**Fig. 3E–G**).

**Figure 3.**
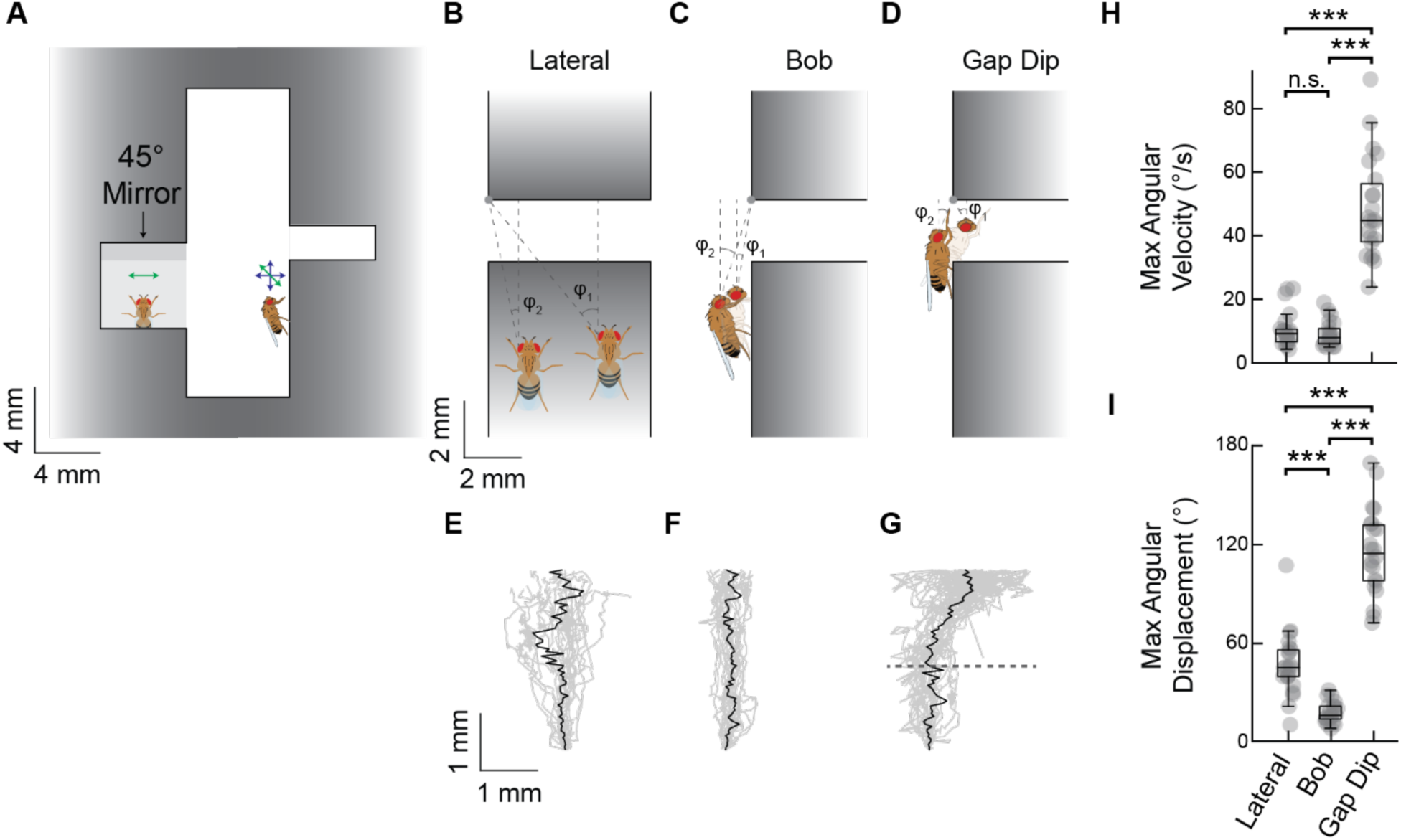
The strongest motion parallax signals during gap encounters come from dipping into the gap. A) Diagram of corridor used to obtain fly position in three dimensions during gap encounters. B–D) Diagrams of three different sources of motion parallax cues during gap encounters: lateral movements in the approach (B), bobs during walking (C), and dips into the gap (D). E–G) Example head trajectories (gray) corresponding to lateral movements (E), bobs during walking (F), and dips into the gap (G), as well as a median head trajectory (black) across all recorded crossings. Dashed line denotes the location of the edge of the gap. H) Maximum angular velocity of the far gap edge relative to the fly head during locomotion from each source of motion parallax cues in (B), (C), and (D). I) Maximum angular displacement of the far gap edge relative to the fly head during locomotion from each source of motion parallax cues in (B), (C), and (D). N = 23 trajectories from 5 flies. Holm-Bonferroni corrected p-values computed using a two-sided Wilcoxon rank sum test. ***p < 0.001.

As a proxy for instantaneous motion parallax information, we computed the maximum angular velocity of the far side of the gap across the fly’s field of view over the course of each individual trajectory (**Fig. 3H**). We computed this based only head translation and ignored body rotations. The maximum angular velocity from motion of the head into the gap was approximately an order of magnitude larger than that generated by either of the two other sources (**Fig. 3H**), reminiscent of findings in mice^35^. In addition to using instantaneous velocity to generate parallax signals, flies could in principle also integrate angular signals over time and thus be sensitive to total angular displacements. To estimate the upper limit of such a strategy, we computed the largest angular displacement of the far side of the gap over the course of each individual trajectory (**Fig. 3I**). The motion of the head into the gap yielded by far the highest angular displacement as in the case of angular velocity, but the angular displacement yielded by lateral movements was significantly higher than that yielded by bobbing movements (**Fig. 3I**). This was because lateral displacements were larger than the head bob displacements, even though the speed of these movements were similar. Altogether, these results suggest that flies may be gathering high fidelity parallax information upon entering the gap.

If parallax information that leads the fly to cross or not is primarily gathered within the gap, we hypothesized that flies would not modulate their locomotor behavior in the approach to the gap. We therefore asked whether it was possible to use the behavior approaching a gap to predict the gap encounter behavior. Neither linear nor non-linear classifiers could predict gap encounter behavior from locomotor behavior only in the approach (**Fig. S3A**). In fact, independent of the gap size and encounter behavior, flies exhibited similar approaches to gaps, in which on average they tended to walk at a roughly constant speed for approximately two seconds while approaching the gap and accelerate slightly in the final half second before reaching it (**Fig. S3B**). This similarity in approach further suggests that flies primarily rely on parallax information gathered within the gap.

### Targeted screen of visual neurons identifies neurons necessary for distance estimation

The visual system of *Drosophila* houses a host of feature-selective neurons projecting to the central brain, some of which are downstream of T4 and T5^52–57^. To find feature-selective visual neurons involved in distance estimation, we performed a silencing screen of visual projection neurons, while also targeting neurons previously implicated in distance estimation^42,44^ (**Fig. 4A**). In our high-throughput rig, we silenced 40 different targeted classes of visual neuron using shibire^ts^ ^47^ and examined the effects on the gap crossing behavior.

**Figure 3. Targeted Figure 4.**
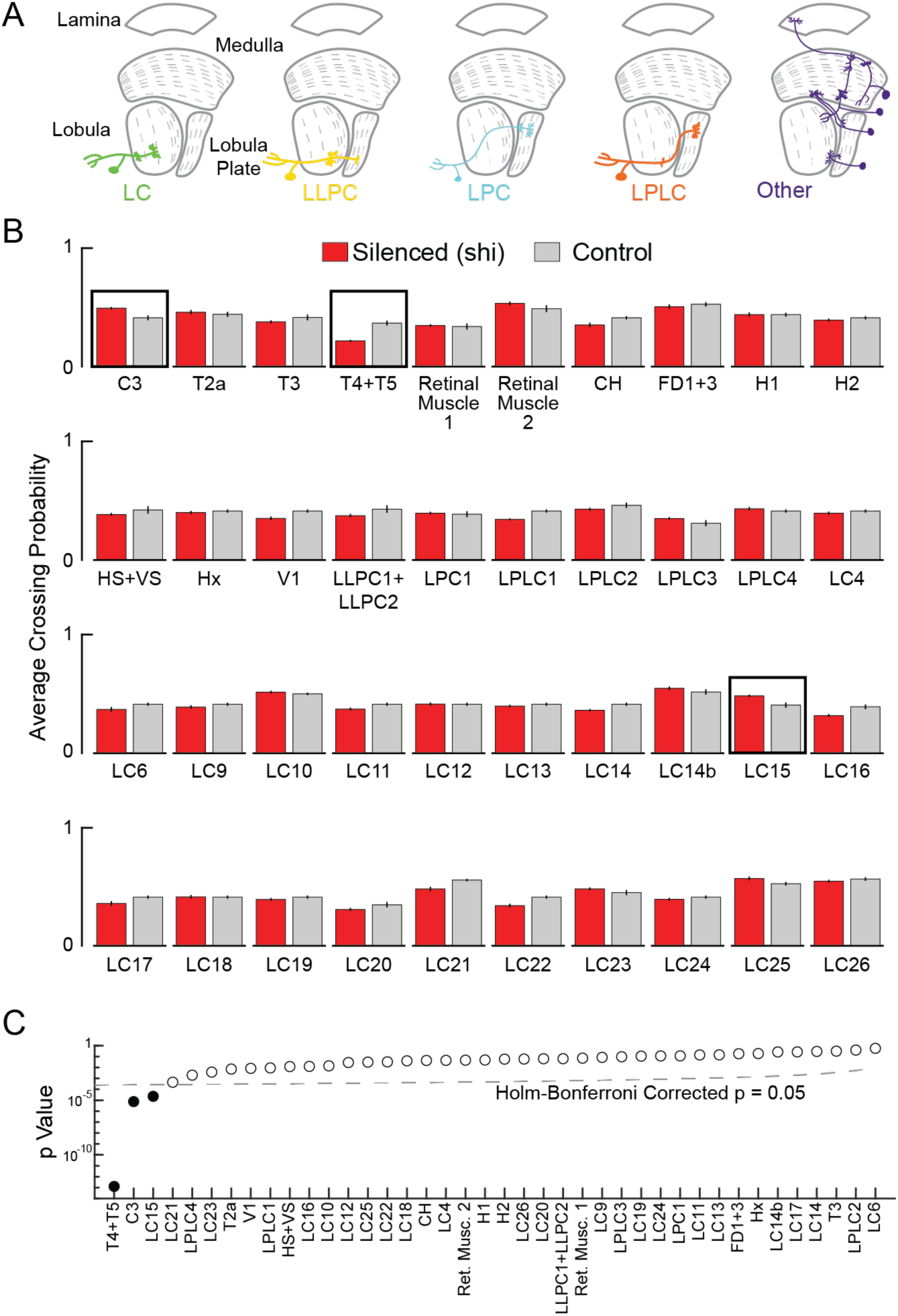
Targeted screen of visual neurons identified neurons necessary for distance estimation. A) Diagrams of the optic neuropils and the neurons tested in the screen. B) Summary of screen data, showing average crossing probability across all gap sizes for all neurons targeted in the screen (silenced neurons in red, X-Gal4 > shi; most conservative control in gray, X-Gal4 > +, empty-Gal4 > shi, or the synthetic control). Error bars show one standard error of the mean. Black boxes denote p<0.05 using Holm-Bonferroni-corrected significance computed from values in the entire crossing probability curves shown in **Fig. S4A** (see **Methods**). N = 12 to 21 flies per genotype. C) Sorted Holm-Bonferroni-corrected p-values for each experiment silencing different neurons (see **Methods** for statistical testing).

Using a conservative statistical testing scheme corrected for multiple comparisons (see **Methods**), we identified three hits from our screen: T4 and T5, C3, and LC15 (**Fig. 4B**). Our T4 and T5 results were described earlier (**Fig. 2**). The second most prominent hit was the neuron C3, a GABAergic medullar neuron that provides feedback to peripheral visual neurons^58^ and was previously implicated in distance perception in gap crossing^42^. Our third hit was the neuron LC15, a visual projection neuron that responds to moving bars in a speed- and direction-insensitive manner^55,59–61^. The effect of silencing C3 and LC15 was more modest than that of silencing T4 and T5 (**Fig. 4B, C**) and shifted the gap crossing frequency curve toward larger gaps (**Fig. 4B, S4A**), consistent with prior findings of silencing C3^42^. Unlike in the neurons T4 and T5, constitutively activating LC15 by expressing the channel dTRPA1 did not result in any change in gap crossing behavior (**Fig. S4B**), suggesting that LC15’s contributions to distance estimation may be more complicated than those of T4 and T5.

### LC15 responds to difference between foreground and background velocities

Because T4 and T5 have well-explored tuning properties^15,46,62–69^ and because C3 neurons are so peripheral in the visual system, we chose to focus our investigations on LC15. LC15 is a feature-selective visual projection neuron that has measured responses to some stimuli^55,59–61^, but not in the context of distance estimation. We therefore decided to characterize its properties to see if they contain the signatures of distance estimation. To do so, we displayed panoramic visual stimuli using a custom rig^70^ while recording calcium responses in LC15 neurons with GCaMP6f^71^ using a two-photon microscope (**Fig. 5A**). LC15 has its dendrites in the lobula and its axon terminals in PVLP (**Fig. 5B, C**).

**Figure 5.**
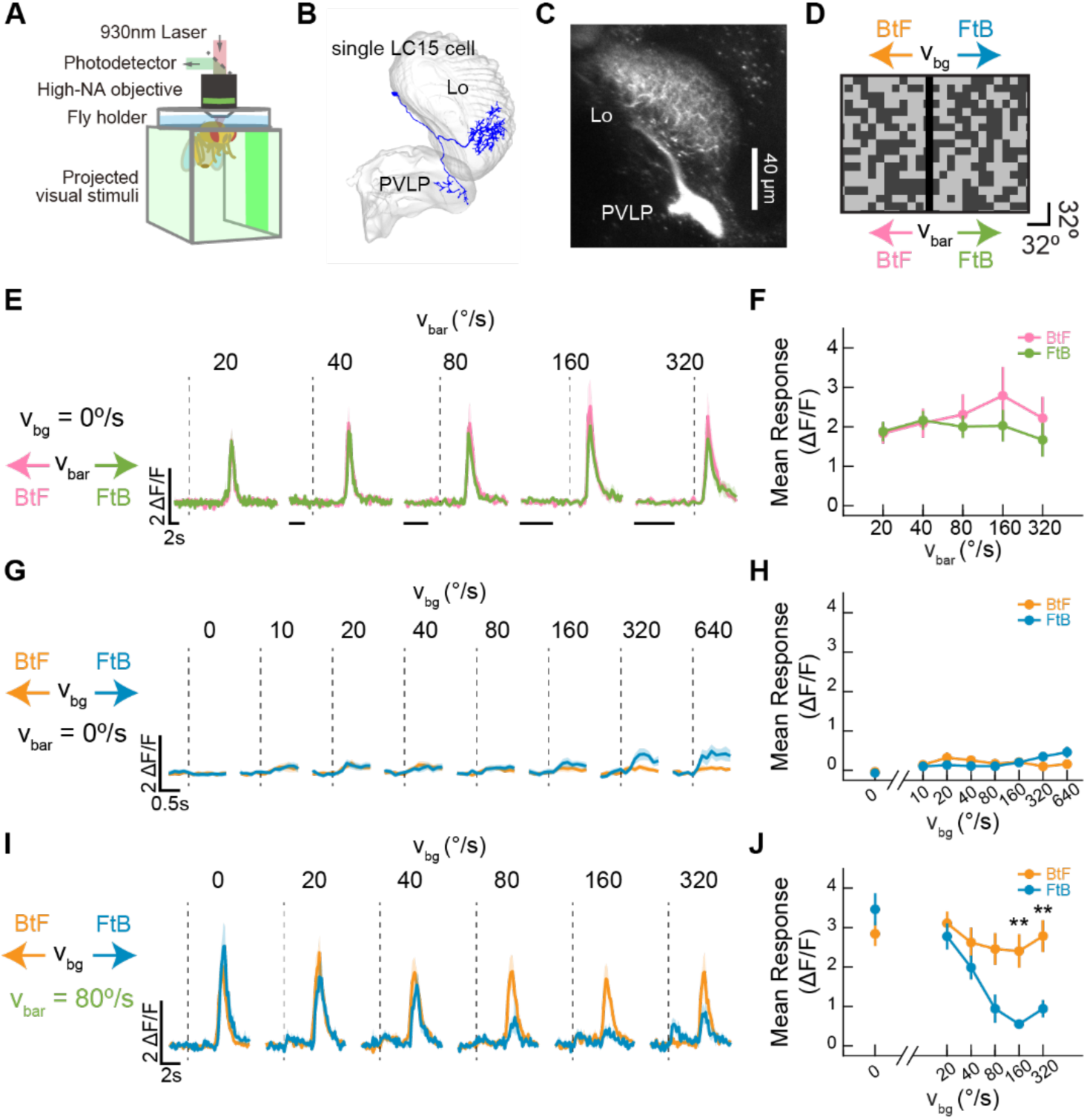
LC15 is relative-direction selective. A) Diagram of two-photon microscopy setup used for *in vivo* calcium recordings during visual stimulation. B) LC15 neurons receive inputs in the lobula (Lo) and project to the posterior ventrolateral protocerebrum (PVLP). C) Two-photon image of LC15 neurons in the optic lobe (LC15 > GCaMP6f). D) Visual stimulus: a random 8° checkerboard background with 50% contrast and a vertical, black 8° bar can move in various combinations of directions (front-to-back or back-to-front) and speeds. E) Mean time traces of LC15 calcium activity in response to the stimulus in (D) with a stationary background and a bar moving front-to-back (FtB) or back-to-front (BtF). F) Time-averaged response across flies for the traces in (E). Error bars represent 1 SEM. G) Mean time traces of LC15 calcium activity in response to the stimulus in (D) with a background moving front-to-back (FtB) or back-to-front (BtF) and no bar. H) Time-averaged response across flies for the traces in (G). I) Mean time traces of LC15 calcium activity in response to the stimulus in (D) with a background moving front-to-back (FtB) or back-to-front (BtF) and a bar moving front-to-back (FtB) at 80°/s. J) Time-averaged response across flies for the traces in (I). N = 12 flies. All error bars and shaded areas represent one standard error of the mean. Holm-Bonferroni corrected p-values computed using a student’s *t*-test. **p < 0.01.

We first presented flies with 8°-wide dark bars moving at various speeds in either front-to-back or back-to-front directions over a random, stationary checkerboard background with 50% contrast and 8° square checks (**Fig. 5D**). LC15 responded roughly equally to front-to-back and back-to-front moving bars and did not show strong speed tuning (**Fig. 5E, F**), consistent with prior measurements^59–61^. We also presented flies the same background pattern moving at different speeds in either the front-to-back or back-to-front directions (**Fig. 5D**). Similar to LC15’s response to bar motion, LC15 responded roughly equivalently to front-to-back and back-to-front wide-field motion and was not speed tuned, but the responses to wide-field motion were far smaller than the responses to moving bars (**Fig. 5G, H**), properties consistent with prior measurements^59–61^.

Previous work had shown that, like many LC neurons, LC15 responses to moving bars are suppressed by background motion^60^. In the context of motion parallax, this sort of suppression offers an opportunity to confer information about object distances. We therefore set out to characterize LC15 responses to relative motion of the bar and the background. To do so, we showed the flies bars moving at 80°/s front-to-back while the checkerboard background moved at various speeds in either the same or the opposite direction of the bar (**Fig. 5D**). Interestingly, the direction of the background strongly modulated the suppression of the bar response (**Fig. 5I, J**). When the bar and background moved in the same direction, the bar response was strongly suppressed, especially when they moved at similar speeds. However, when the background moved in the opposite direction of the bar, there was virtually no suppression. This means that, although LC15 does not exhibit direction selective responses to either the bar or the background individually, it exhibits strongly relative-direction selective responses that depend on the joint direction of the bar and background motion.

### Relative motion is informative of distance

Because LC15 exhibited relative-direction selectivity for joint foreground and background motion, we first asked how informative the relative motion between foreground and background can be about distance. In a motion parallax signal generated by an observer’s translational movement, the relative distance from the observer to two features in the visual field (such as an object and a background) is inversely proportional to the relative retinal speeds of the features (**Fig. 6A**). Therefore, a motion parallax-based distance detector should in principle be tuned to respond primarily to the ratio of the velocities of two features in the visual field, independent of their individual velocities.

**Figure 6.**
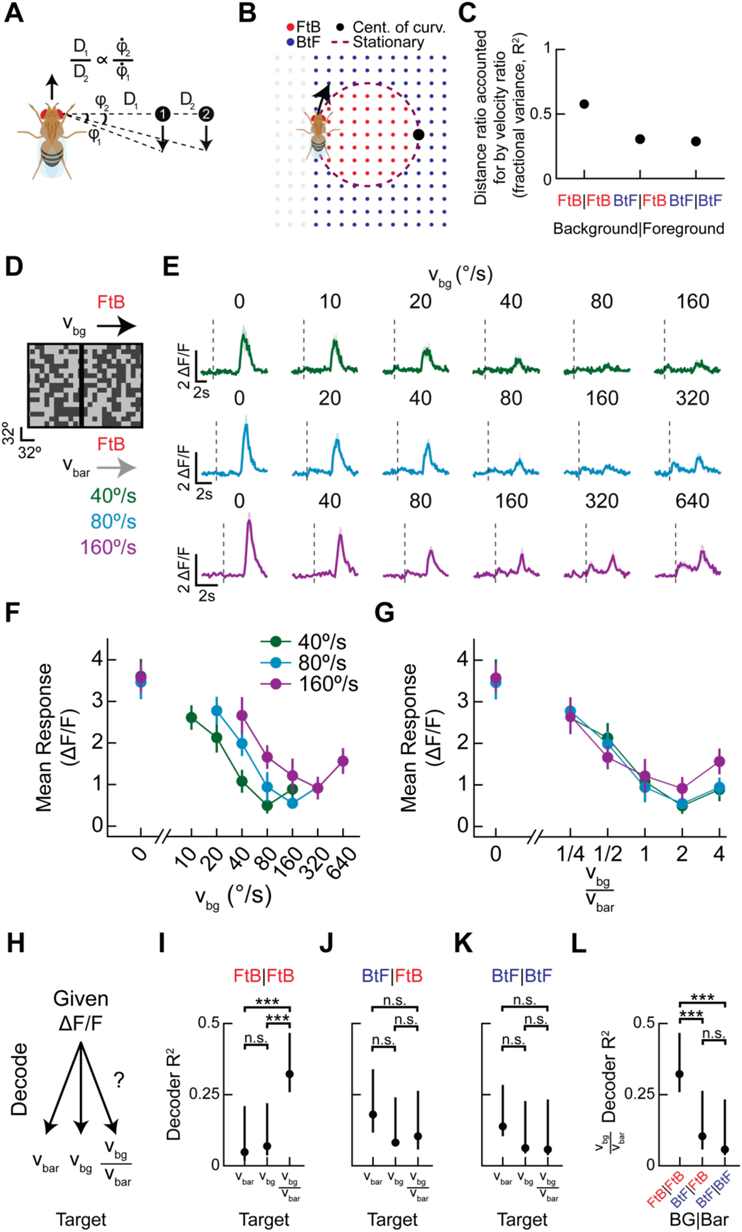
LC15 exhibits parallax-like tuning to relative motion. A) Diagram showing geometric parametrization of motion parallax. B) Diagram showing the direction of motion in the fly’s frame of reference of stationary points as the fly follows the trajectory in black. Depending on the angle from the fly and the fly’s curvature, points nearby move front-to-back (FtB), points more distant than the center of curvature (black dot) move back-to-front (BtF), and a subset of points are stationary on the eye (dashed purple). C) In a simulation of fly walking trajectories and object distances, for each combination of front-to-back (FtB) and back-to-front (BtF) motion of a closer object and further object (assigned foreground and background objects), we computed how much variance in the distance ratio could be accounted for by the velocity ratio (see **Methods**). Error bars of one SEM in our simulation are smaller than the drawn points. D) Visual stimulus in which a vertical bar moves front-to-back at one of three velocities while the random checkerboard background moves front-to-back at several velocities. Bar and background were as in Figure 5. E) Mean time traces of LC15 calcium activity in response to the moving bar and background as described in (D). Shaded area shows ±1 SEM. N = 12 flies F, G) Time averaged response across flies of the traces in (E) plotted against the background velocity (F) and plotted against the ratio of background and bar velocities (G). Error bars represent one standard error of the mean. H) A non-parametric decoder was trained to predict the velocity of the bar, the velocity of the background, and the ratio of the velocities, given the responses measured in LC15. I) The variance predicted by the decoder of neural responses for bar velocity, background velocity, and the ratio of the two velocities. In this case, both the bar and background moved front-to-back. J) As in (I) but for stimuli in which the bar moves front-to-back and the background moves back-to-front. K) As in (I) but for stimuli in which the bar and background both move back-to-front. L) For each of the three measured combinations of front-to-back and back-to-front bar and background directions, the variance predicted by the decoder for the ratio of the bar and background velocities. Error bars in (I)–(L) represent the 5^th^ and 95^th^ percentiles of bootstrap resamplings (see **Methods**). p-values computed via bootstrapping. ***p < 0.001.

Importantly, this correspondence between relative distance from the observer and relative retinal speed only holds in the case of purely translational motion. When an observer rotates while translating, this correspondence is no longer true for all pairs of features. In this case, the speed and even the direction of motion of a feature depends both on its distance from the observer and on the radius of curvature of the observer trajectory (**Fig. 6B**). Objects further than the center of curvature tend to move back-to-front across the retina, even when the fly is walking forward^15^. For any given trajectory, the relative distance and relative retinal speed can be computed geometrically. To better understand how this geometry affects motion parallax encoding, we quantified the relationship between the retinal speed ratio and the object distance ratio under a variety of conditions. To do this, we simulated an ensemble of trajectories and object/background distances, with parameters approximating typical walking trajectories^72,73^ (see **Methods, Fig. S6A–E**). We next asked how much variance in the relative object/background distances could be predicted from knowing the ratio of the retinal velocities (see **Methods**). When both the object and the background were moving front-to-back across the retina, the retinal velocity ratio predicted the relative distance well (**Fig. 6C**). This situation is one in which objects are closer than the center of curvature of the fly trajectory, which can happen when flies move in relatively straight lines, as they often do^72^. In other cases, in which one or both features are farther than the center of curvature during walking, the ratio of the retinal velocities accounts for a substantially smaller fraction of the variance in relative distance (**Fig. 6C**). Overall, these results show that when features are closer than an observer’s center of curvature, relative motion remains highly informative of distance, but when features are further away than the center of curvature, relative motion is less informative about distance. The informative situation corresponds specifically to a stimulus in our paradigm in which both foreground and background patterns move front-to-back across the retina.

### LC15 exhibits parallax-like tuning to relative motion

Given LC15’s relative-direction selectivity and its disruption of gap crossing behavior when silenced, we wanted to test whether LC15 encoded motion parallax information in a manner consistent with this geometrical model for motion parallax. To investigate this, we presented the same bar and random checkerboard background as before, but now with three bar speeds and a suite of background speeds, all in the front-to-back direction (**Fig. 6D, E**). For all three tested bar speeds, the response was suppressed by some background velocities, with peak suppression when the background speed was faster than the bar speed. Strikingly, the curves collapsed onto one another when plotting the response against the ratio of the background and bar velocity (**Fig. 6G**).

The collapse of the curves means that LC15 neurons responded approximately the same way, not to the bar velocity or to the background velocity, but to their ratio, over a factor of four in bar speeds. We also tested other combinations of bar and background directions, and this collapse did not occur for the other combinations of bar and background directions that are realizable by a forward walking fly (**Fig. S6F–K**). In those other direction combinations, the geometry indicates that the ratio of the velocities does not correspond as strongly to the ratio of distances (**Fig. 6C**). These results show that LC15 responds to the velocity ratio in the instance where that ratio is most predictive of the distance ratio but not in the instances where that ratio is more weakly predictive of the distance ratio. Thus, the LC15 responses to front-to-back moving stimuli suggest that it is tuned to the relative distance of the object and background.

To quantify the qualitative matching of the responses when plotted against bar-background velocity ratios, we tested whether we could use LC15 responses to decode different stimulus quantities, including the bar velocity, background velocity, and their ratio (**Fig. 6H**). To do this, we fit three non-parametric decoders to the recorded neural responses in each condition (see **Methods**): one to predict the bar velocity, one to predict the background velocity, and one to predict the ratio of the background to bar velocities. If LC15 responses best represented the ratio of the visual velocities, then the decoder trained to predict the ratio of the velocities should perform the best. Indeed, the decoder to predict the ratio of the velocities from the LC15 response performed significantly better than the other two decoders in the case of front-to-back motion of both the background and bar (**Fig. 6I**). This indicates that LC15 signals best encode the ratio of the two velocities, and thus relative distance. Moreover, the velocity ratio was only more decodable in the configuration in which both bar and background moved front-to-back (**Fig. 6I-K**), and the velocity ratio was significantly more decodable in the front-to-back configuration than in either of the other two directional configurations (**Fig. 6L**). Notably, the front-to-back configuration is also the configuration in which the ratio of the velocities is most informative about relative distances (**Fig. 6C**). Altogether, LC15’s identification in the behavioral screen, its observed neural signals, and its alignment with the geometrical model for the usefulness of parallax signals all provide compelling evidence that LC15 encodes motion parallax cues in a coherent, contextual manner.

## Discussion

Using our high-throughput gap crossing assay, we genetically silenced a large set of neurons in the fly visual system and identified multiple neurons involved in the perception of visual distance. Our assay found that binocular vision was dispensable for judging gap distance in this assay, consistent with prior findings^25^. Our screen identified one neuron class identified in a prior study^42^, strongly implicated the primary motion detectors in distance perception, and discovered a novel feature-detecting neuron that encodes motion parallax cues.

### Local motion detectors, T4 and T5, are required for distance estimation

Our discovery that the primary visual motion detectors, T4 and T5, are necessary for distance estimation in gap crossing (**Fig. 2C, D**) provides further evidence that motion parallax serves as the primary visual cue used by flies in this paradigm^25^ and adds to the behaviors that rely on signals from T4 and T5^15,46,74,75^. In primates^33^ and mice^35^, causal links have been found between specific cortical areas and distance estimation, but here we were able to link distance perception to a single pair of neuron types. Identifying individual feature-selective neurons required for distance estimation provides a clear path for investigating the distance perception circuit: search in the circuitry downstream of T4 and T5 for parallax signals and for distance signals deeper in the brain. Our screen did not identify hits in neurons directly downstream of T4 and T5, though we did test several, including LPLC1 and LPLC2, which are loom-selective^59,76,77^ and therefore conceivably involved in judging distance. There remain a small set of visual projection neurons downstream of T4 and T5 that we did not test because we lacked drivers, and there is also a large set of tangential cells that could be involved^56,78^. It may be difficult to identify these neurons, just as it has been difficult to identify the circuitry directly downstream of T4 and T5 involved in orientation control, one of their other primary functions.

### Non-canonical parallax response properties of LC15 neurons

Silencing LC15 in our behavioral screen led to a modest change in gap crossing behavior (**Fig. 4B, S4**), similar to the change we and a previous study^42^ observed when silencing C3 (**Fig 4B**). However, LC15 is deeper in the visual processing circuits than C3 and is a feature-detecting neuron. Our characterization of LC15 shows how its neural responses can encode relative distance between objects and backgrounds (**Fig. 6**). Its response properties are similar to measurements of distance- and disparity-tuned neurons reported in primates^17,23,24,33,40^ in that differing ratios of object distances correspond to differing neural responses. LC15 differs from the reported primate neurons in two key respects. First, it was identified through the behavioral consequences of its silencing. The silencing experiments tie the neural activity of LC15 neurons directly to distance-judgment behavior, which has been difficult in primate models. Second, LC15 is not direction selective for objects or wide-field motion (**Fig. 5F, H**). Instead, it shows relative-direction selectivity in its gain, so that it is more suppressed when the background and foreground both move front-to-back than when they move in opposite directions (**Fig. 5J**). It is unclear how such relative-direction selectivity is implemented. One possibility is that LC15 integrates inputs corresponding to object- and widefield-motion from its upstream partners, which in turn receive direction selective inputs from T4 and T5. The most plausible candidate neurons are TmY9q, TmY9q^⊥^, and Li27, identified in the fly connectome^79–81^.

LC15 neurons display a rich set of response properties (**Fig. 6**) that add to the diversity of feature-selectivity found in projection neurons in the *Drosophila* lobula. Prior work on visual projection neurons has shown tuning to object size, polarity, direction, and speed^54,55,59–61,76,82–85^, as well as to growth in feature size (looming)^77,86,87^. Responses in various feature detectors have been shown to be suppressed during wide-field motion^60,61,76^. This work shows that feature detectors can respond in sophisticated ways to the joint speeds and directions of visual patterns, in ways that are tuned to the three-dimensional organization of the world. This research suggests that other feature detectors in the fly eye may possess similarly rich response properties that can be elucidated by tying them to specific visual behaviors.

### Spatial understanding of motion parallax cue relevance

Previous studies have predominantly investigated motion parallax in animals that fixate using eye movements^17,23,24,33,40^ or compensatory head movements^22^. Such fixation and the resulting geometry lead to depth-sign ambiguity^22^, the phenomenon in which it is ambiguous whether an object is closer or further than the fixation point. Depth-sign ambiguity has been shown to be resolved in primates by integrating extra-retinal signals, which can indicate direction of the observer’s translation^17^.

Through our simulation of parallax signals generated by observer translation and rotation (**Fig. S6A–E**), we showed that there exists depth-sign ambiguity around the center of rotation, which is geometrically similar to a fixation point in classical vertebrate studies. However, the ambiguity is resolved because flies primarily walk forward^72,73^, so that front-to-back motion across the retina is typically closer than back-to-front motion (**Fig. 6B**). Within this curved-walking framework, certain regimes of retinal motion exhibit stronger correspondence between motion parallax cues and feature distances (**Fig. 6C**), and these regimes correspond to those in which LC15 exhibits tuning for the informative ratio of bar and background velocities (**Fig. 6L**). By contextualizing parallax information in a geometrical simulation, we have shown how the curvature of a moving observer’s trajectory can influence the usefulness of parallax cues for determining distance, consistent with recent work in primates^88^.

### Breadth of response properties encoding motion parallax cues

Although studies have shown animals to respond in a non-direction selective manner to some forms of parallax stimuli^15,22^, to our knowledge, all previously identified disparity- and distance-selective parallax neurons have been direction selective^17,23,24,33,40^. This is likely in part because depth-sign disambiguation is likely to rely on direction selective information^17^. In contrast, this study demonstrates that neurons may exhibit parallax-tuning despite lacking canonical directional properties. We show that both direction-selective T4 and T5 neurons and non-direction-selective LC15 neurons contribute to distance estimation in fly gap crossing behaviors. This suggests that non-directional neurons could contribute to distance estimation in other visual systems as well, potentially in combination with canonical distance signals in direction selective neurons. Overall, our results highlight a sophisticated fly feature detector and the breadth of feature detector properties that can convey information about relative object distance.

## Contributions

JS, EW, RT, and DAC conceived of the behavioral experiments. JS and EW obtained and analyzed behavioral data for the behavioral screen. EW and BA obtained and analyzed neural imaging data. KM, NM, G Santana, EW, and JS obtained and analyzed behavioral data for visual cue involvement. BZ, G Sager, and JS developed classification models. EW, JS, BA, KM, and CG created figure panels. RT, EW, and JS created fly lines used in experiments. JS wrote the first draft of the paper; it was edited by JS, RT, EW, and DAC.

## Acknowledgments

We thank Claude Desplan for helpful advice on visual projection neuron driver lines, as well as Alec Sheffield, Kevin Chen, Clayton Barnes, and Emma Flor for helpful comments on the manuscript. We thank Mahmut Demir and Hany Dweck for invaluable advice on the experimental set up. JS was funded by a Ford Foundation graduate fellowship. G Sager, KM, and CG were funded by NSF GRFs. BA was funded by a NSERC graduate fellowship. RT was funded by a Takenaka Foundation fellowship. DAC and this project were funded by NIH R01 EY026555.

## Figures

**Supp. Figure S1.**
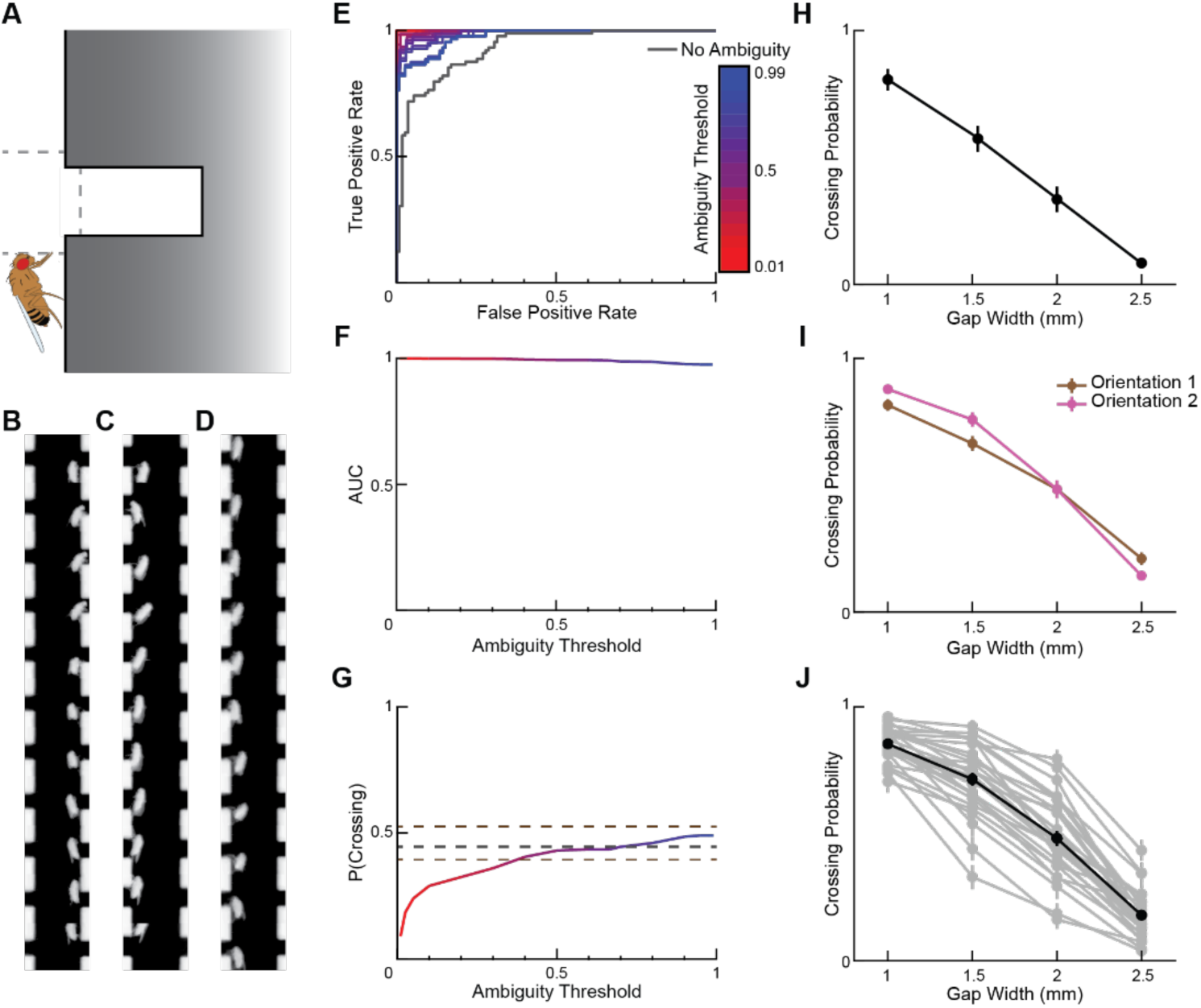
A neural network classified gap encounter behavior. A) Diagram of how gap encounter behaviors were triggered based on the fly location. Once the head passed boundaries surrounding the gap (dashed gray lines, 0.5 mm away from the gap edges), the fly was considered to have moved from one region to another, marking the start of a gap encounter B–D) Examples of image sequences provided to the classifier neural network for training. These sequences were manually annotated as unambiguous crossing (B), unambiguous glass circumvention (C), and ambiguous crossing (D). The full depth of the gap was truncated in the images provided to the classifier. E) A neural network was trained to classify the encounter types (see **Methods**). Receiver operator curve (ROC) for the neural network trained to specifically label gap encounters as crossings or glass circumventions. Each different colored curve represents a different ambiguity threshold set for the network, as described in **Methods**. The ambiguity threshold used for all results in the main figures was 0.5. F) Area under curve (AUC) as a function of ambiguity threshold for the ROCs in (E). G) Probability for the neural network to predict a gap encounter as a crossing as a function of ambiguity threshold. The black dashed line represents the probability of a crossing in the manually annotated training set, and the brown dashed lines represent the upper and lower bounds of the manually annotated training set based on manual annotations on the ambiguity as described in **Methods**. H) Probability of crossing of WT flies for each gap size in a horizontally oriented cassette. N = 9 flies. I) Probability of crossing among WT flies for each gap size in a vertically oriented cassette in orientation 1 (brown) and orientation 2 (pink). N = 28 flies. J) The mean (black) and individual (gray) probability of crossing among WT flies for each gap size in a vertically oriented cassette. N = 28 flies. Error bars represent one standard error of the mean.

**Supp. Figure S2.**
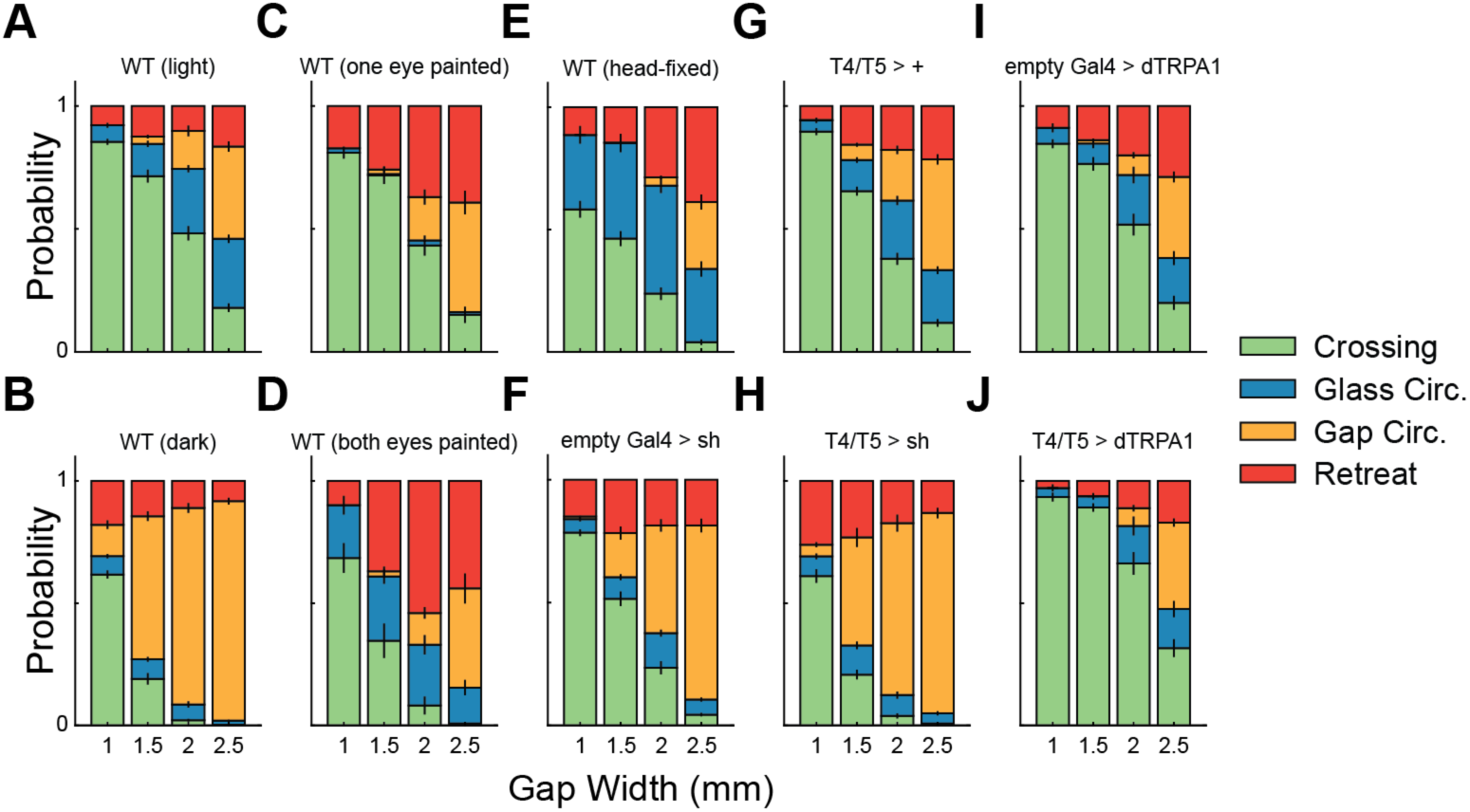
All gap encounter types for flies in the dark, eye-painted flies, and T4 and T5 manipulations. A–J) Probability of each gap encounter event for each gap size among WT flies (A), WT flies in the dark (B), WT flies with one eye painted (C), WT flies with both eyes painted (D), WT flies with head glued to abdomen (E), a neuronal silencing parental control (F), a T4 and T5 driver parental control (G), T4 and T5 silenced flies (H), a neuronal activation parental control (I), and T4 and T5 activated flies (J). N = 12 to 28 flies per genotype. Error bars represent one standard error of the mean.

**Supp. Figure S3.**
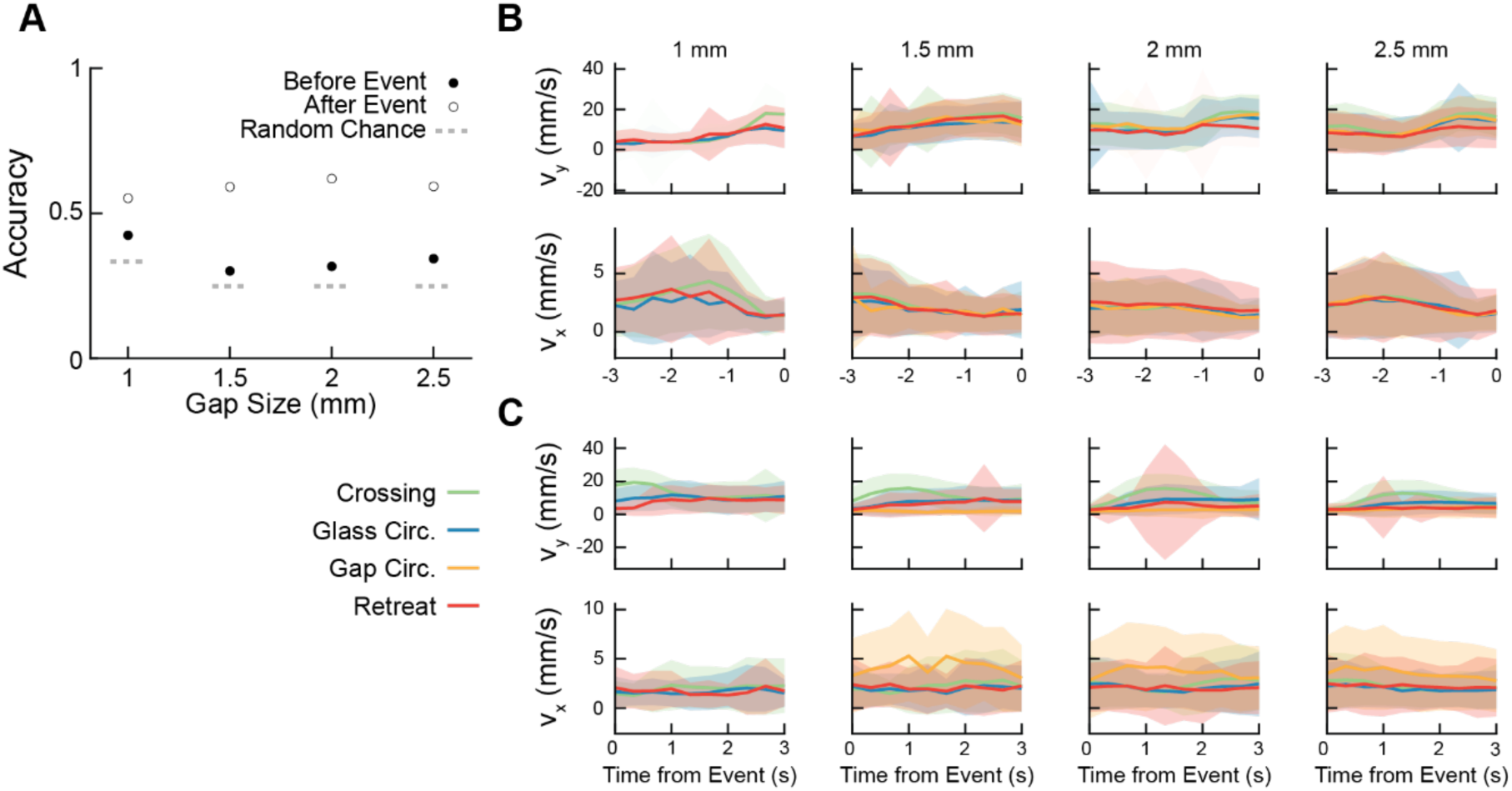
Flies approach gaps in a stereotyped manner independent of gap size and gap encounter behavior. A) Two non-linear classifiers were trained on trajectories to try to predict the gap encounter behavior at each gap size: one using trajectory information preceding the encounter (filled circles) and one using trajectory information following the encounter (open circles). The classifier trained on information preceding the event performs barely better than chance (gray), suggesting that flies approach gaps in a stereotyped manner independent of gap encounter behavior and gap size. The classifier trained on information following the encounter serves as a positive control to ensure that such a non-linear classifier is capable of classifying the gap encounter behavior given sufficient relevant information. B) Average vertical and horizontal velocity of fly centroids in the three seconds preceding a gap encounter for each gap size and each behavior. C) Average vertical and horizontal velocity of fly centroids in the three seconds following a gap encounter for each gap size and each behavior. Shaded areas represent one standard error of the mean. N = 28 flies.

**Supp. Figure S4.**
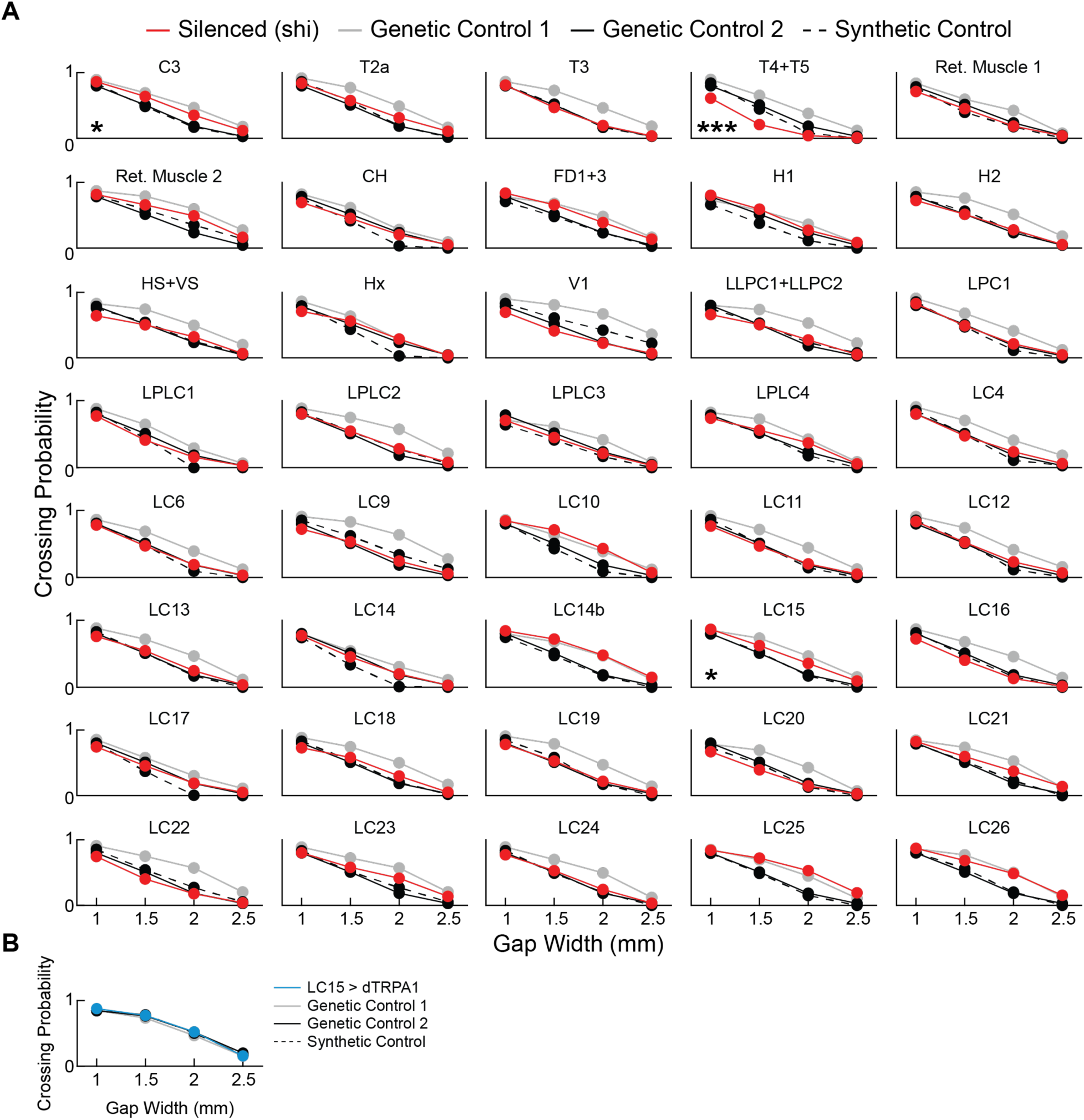
Full crossing curves from the targeted silencing of neurons in the screen. A) Probability of crossing for each gap size. Each panel shows the silenced neuron (red, X-Gal4 > shi), two parental controls (gray, X-Gal4 > + and solid black, empty Gal4 > shi), and a synthetic control (dashed black) that represents a linear additive model of the two parental controls (see **Methods**). Neurons in the screen were only counted as hits if the silencing curve was statistically significantly different from both parental controls and from the synthetic control post Holm-Bonferroni correction, providing a conservative test for observed effects. B) Probability of crossing for each gap size for flies with constitutively active LC15 (blue, LC15 >dTRPA1), two parental controls (gray, LC15 > + and solid black, empty Gal4 > dTRPA1), and the synthetic control (dashed black). N = 12 to 21 flies per genotype. Error bars show one standard error of the mean but are typically smaller than the dots. Holm-Bonferroni corrected p-values computed using a two-sided Wilcoxon rank sum test for comparisons to parental controls and using a *z*-test for comparisons to the synthetic control. *p < 0.05; ***p < 0.001.

**Supp. Figure S6.**
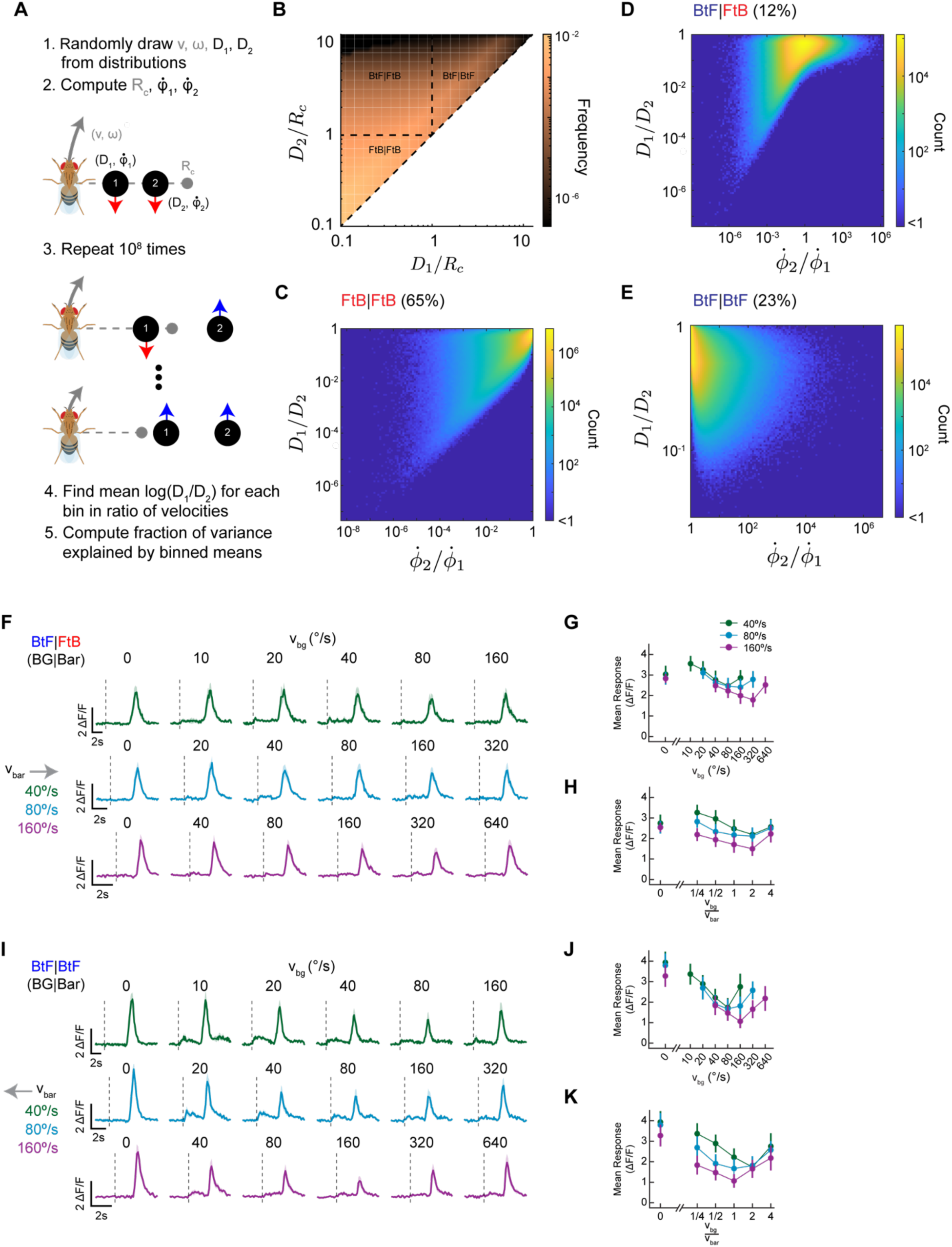
A geometrical simulation shows relationship between angular velocity ratios and distance ratios for agents that both rotate and translate; LC15 calcium traces for different combinations of bar and background velocities. A) Steps involved in computing parallax correspondence in space using a geometrical simulation. For each simulation, the forward walking speed and rotational velocity of the fly were drawn from a distribution of typical speeds (see **Methods**). Likewise, the distance of two features were randomly drawn from a distribution. Combined, these were used to compute the radius of curvature and the retinal velocities of the two features in the fly frame. This process was repeated 10^8^ times to create an ensemble of simulations. B) The simulation distributions were visualized by binning in two dimensions, D_1_/R_c_ and D_2_/R_c_, where R_c_ is the radius of curvature. Different regions of this plot correspond to different directions of motion on the retina, designed by front-to-back (FtB) and back-to-front (BtF) for the far object | near object. C) The distribution of *D*_1_/*D*_2_ and *ϕ̇*_1_/*ϕ̇*_2_ for cases in which both objects were moving front-to-back. This represented 65% of simulated cases in (B). D) As in (C), but for the near object moving front-to-back and the far object moving back-to-front. This represented 12% of simulated cases in (B). E) As in (C), but for the near and far objects both moving back-to-front. This represented 23% of simulated cases in (B). F) Mean time traces of LC15 activity in response to a black bar that moves front-to-back (FtB) at one of three velocities while a half-contrast random checkerboard background moves back-to-front (BtF) at one of several velocities. Bar width was 8° and background check sizes were 8°. G, H) Time averaged response across flies of the traces in (F) plotted against the background velocity (G) and plotted against the ratio of background and bar velocities (H). Error bars represent one standard error of the mean. I) Mean time traces of LC15 activity in response to a black bar that moves back-to-front (FtB) at one of three velocities while a half-contrast random checkerboard background moves back-to-front (BtF) at one of several velocities. Bar width was 8° and background check sizes were 8°. J and K) Time averaged response across flies of the traces in (I) plotted against the background velocity (J) and plotted against the ratio of background and bar velocities (K). Error bars represent one standard error of the mean. N = 12 flies for panels F–K.

**Supplemental Movie 1.** Movie of flies walking in vertically oriented corridors during experiment. Three trials are shown with two flips between them. Computed fly positions are shown in red.

## Methods

### Code and data availability

Raw imaging data is available on DANDI here: [link on publication]. Processed behavioral and imaging data traces, along with code to make figures, is available on Dryad and Github here: [links on publication]. And code for the distance simulation is available on Github here: [link on publication].

### Fly Strains

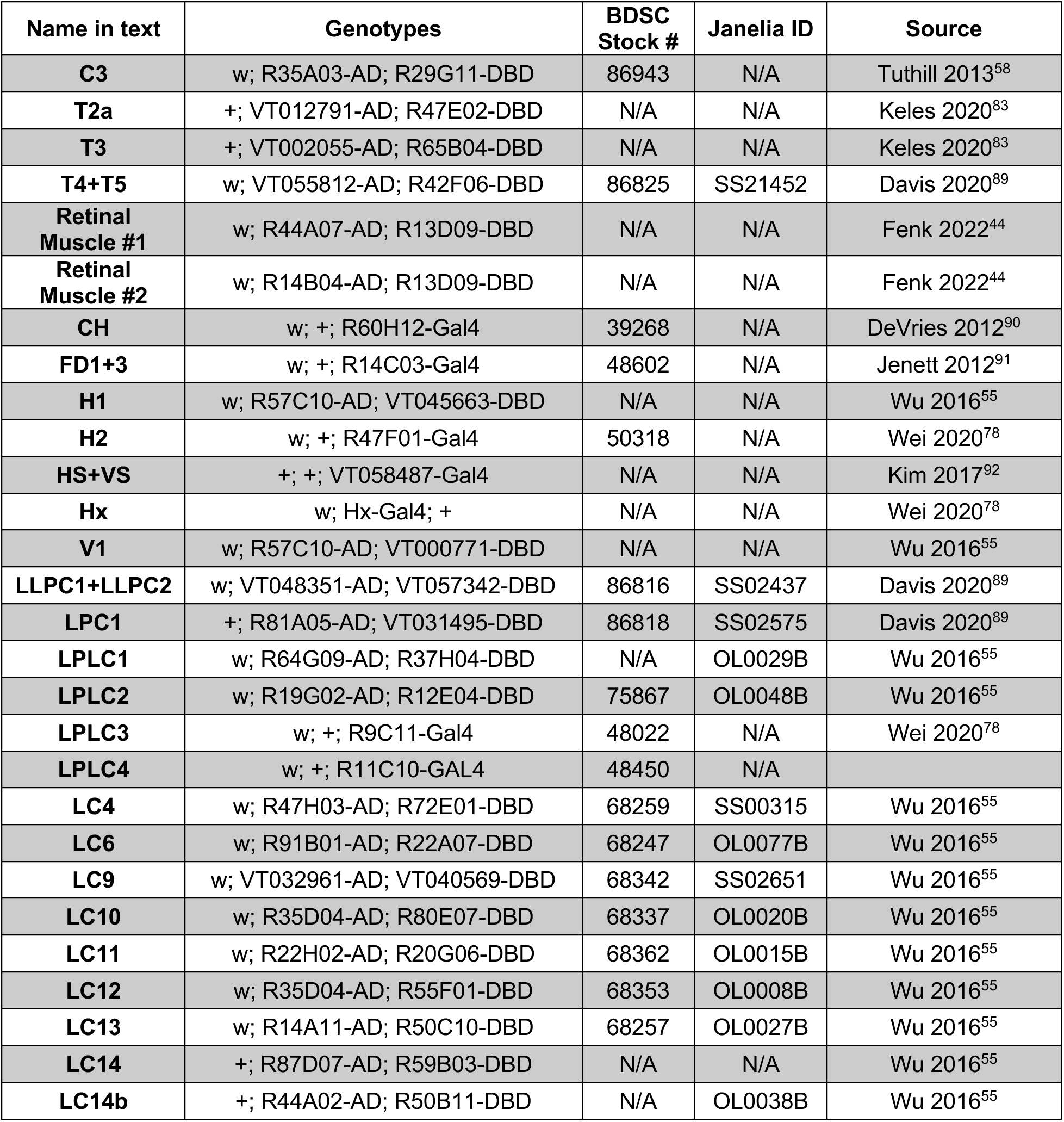
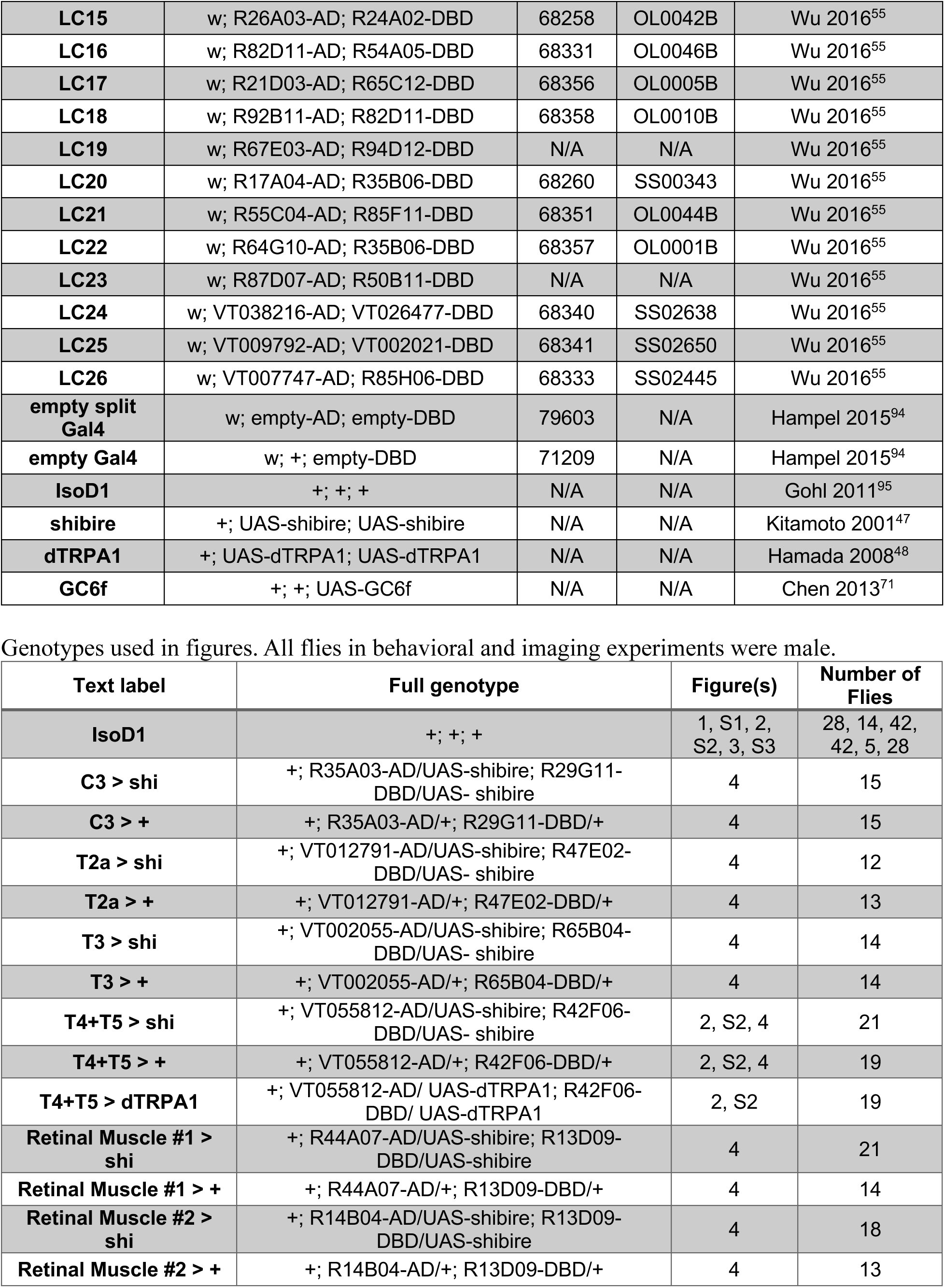
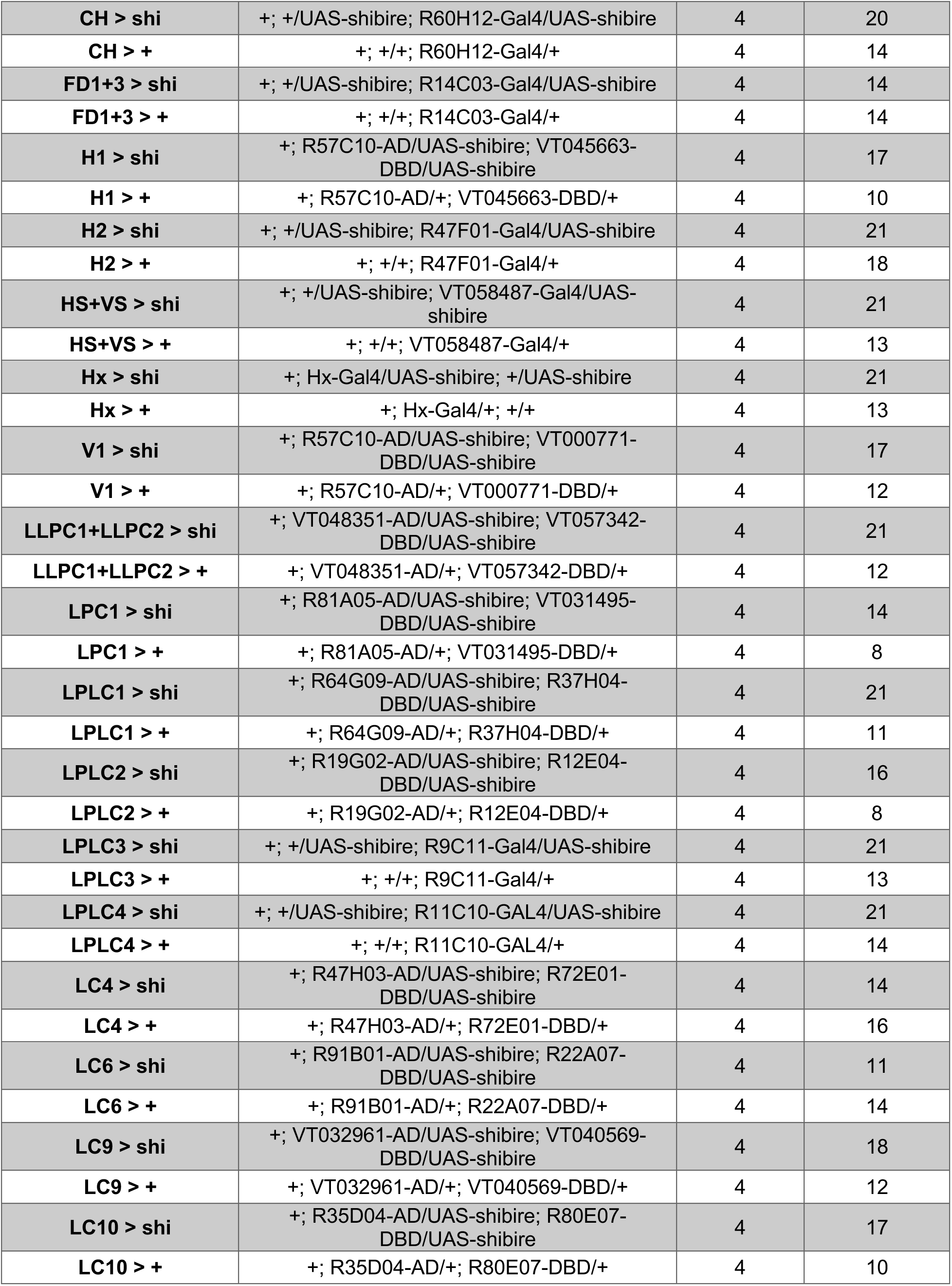
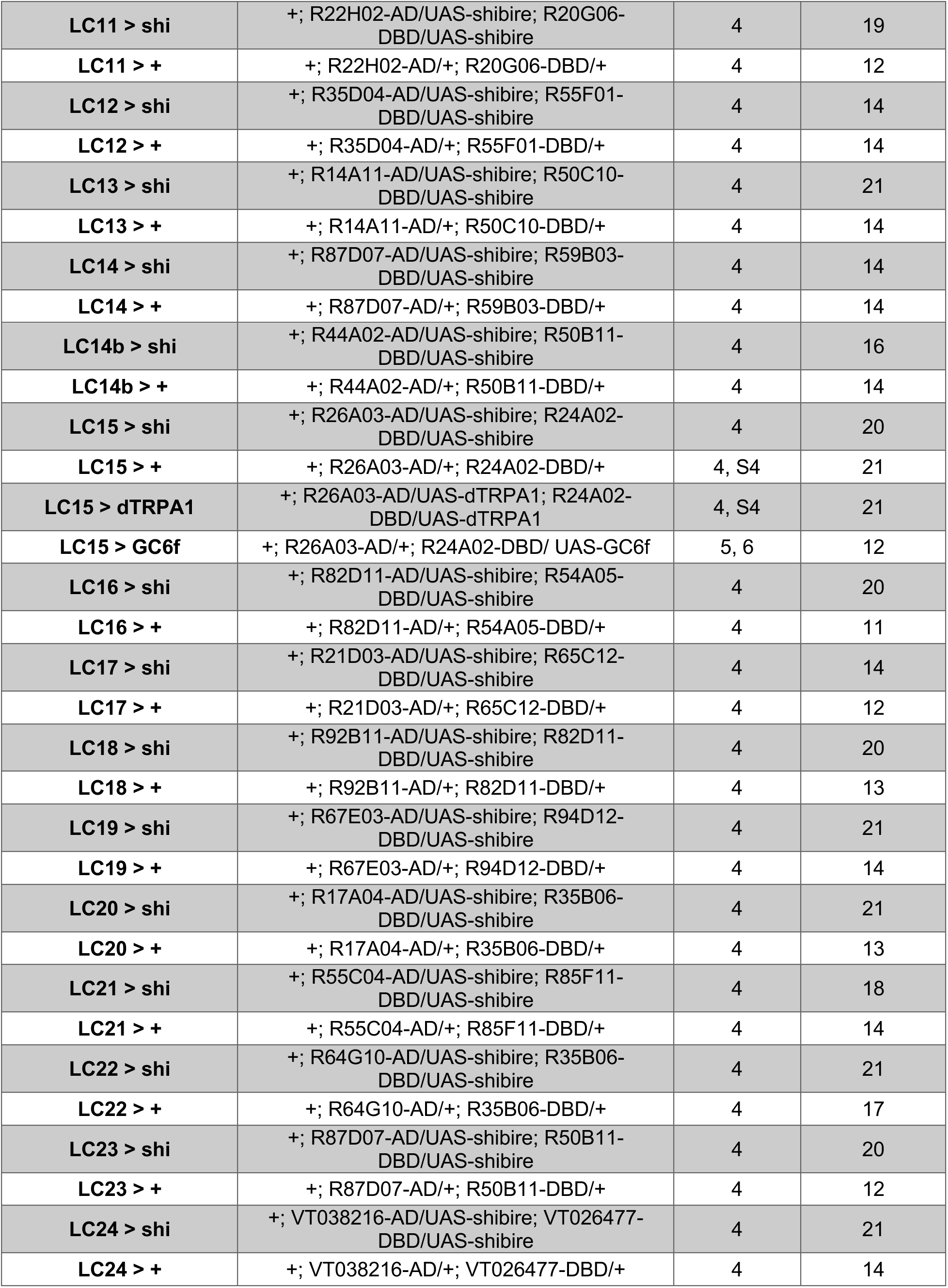
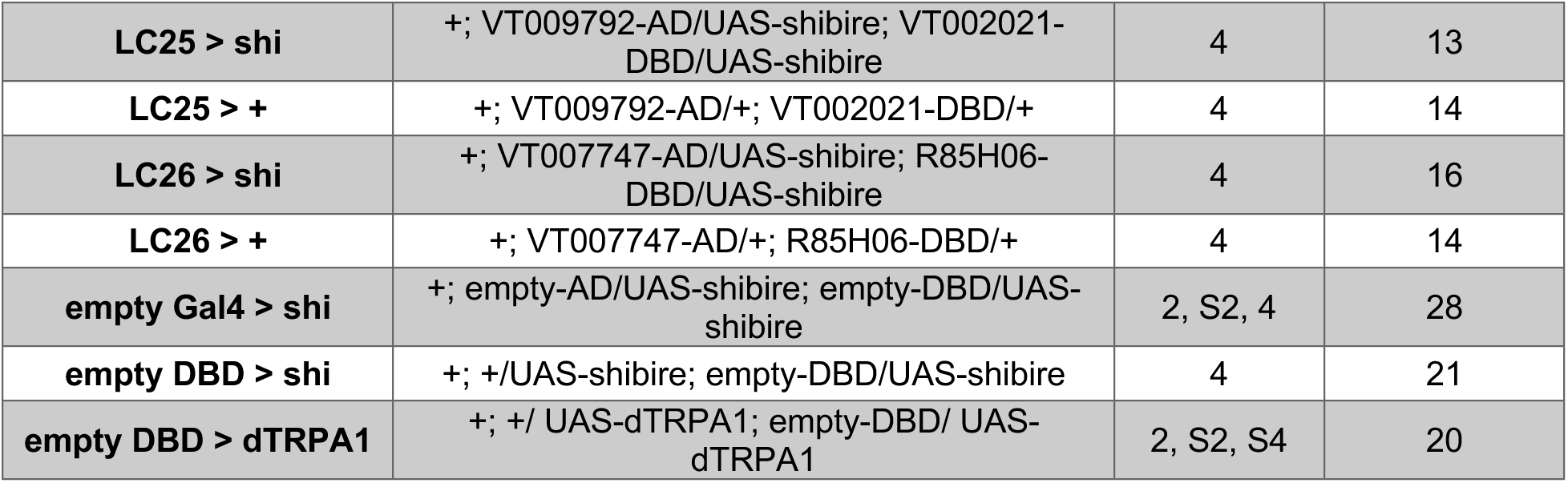

### Fly rearing

All flies were reared in incubators in a room maintained at 50% relative humidity. We used exclusively male flies for all behavioral and imaging experiments in this research. Flies used in behavioral and imaging experiments were reared at 20 °C and 25 °C, respectively, and raised on glucose food (recipe D20102, Archon Scientific, Durham, NC, USA). Flies were circadian entrained on 12-hour light-dark cycles and tested for behavior in the three hours after dawn or before dusk. CO_2_ was used to anesthetize flies to stage them at least 24 hours before they were used for behavioral or imaging experiments.

### Behavioral gap crossing assay

Gap crossing cassettes consisted of a laser cut 7"x4"x1/4" ProofGrade Walnut Plywood panel (GlowForge, Seattle, WA, USA); this corridor was found to elicit more robust walking behaviors than various plastics and metals. It was cut new every ∼10 experiments. The corridors were sandwiched between two panes of 7"x4"x1/8" borosilicate glass (part 8476K49, McMaster Carr, Elmhurst, IL, USA) each coated with 0.1 mM berberine chloride (item 00900585, Sigma-Aldrich, Burlington, MA, USA) with Rain-X Original Glass Water Repellent Trigger (Rain-X, Cleveland, OH, USA) used as the solvent. The coating discouraged flies from walking on the glass panes by providing poor physical contact and a bitter taste. The coating was applied following the standard Rain-X application procedure of repeated overlapping circles using a microfiber cloth or paintbrush until no more beaded droplets remained on the glass surface. Unless otherwise stated, each cassette had seven independent corridors that each held one fly and had gap sizes of 1, 1.5, 2, and 2.5 mm. The mirror experiments consisted of gap crossing cassettes with a few different features: each cassette only contained two corridors to allow for higher resolution recording of each fly; each corridor was horizontally asymmetric to accommodate the gap and the mirror (part MRA05-G01, ThorLabs, Newton, NJ, USA); each corridor only had one gap; and the walking surface was covered in Teflon tape (item S-14666, ULine, Pleasant Prairie, WI, USA) to decrease the impact of shadows under the flies in the mirror. The gap crossing cassettes were loaded into a custom-made periodic flipping robot consisting of felt grips rotated by a high-fidelity servomotor (part 6627T53, McMaster Carr, Elmhurst, IL, USA) controlled by an Arduino Uno board. The flipping robot flipped the cassette 180° every eight seconds (slightly longer than the average time it takes a fly to navigate from the bottom of the corridor to the top). The flipping robot was kept in a light isolated box maintained at 34 °C within a room held at ∼50% relative humidity.

Flies were used for behavioral experiments 24-36 hours after staging and within the three-hour windows after circadian dawn or before circadian dusk. When staging flies, we selected male flies and housed them in vials of seven to nine flies. For loading into the gap crossing cassettes, flies were transferred into empty vials and cold-anesthetized in ice water for 90 seconds, after which they were individually loaded into corridors in a cassette. We empirically found that flies exhibit a period lasting ∼5 minutes post-cold-anesthetization during which any exposure to a bitterant (such as berberine chloride) caused them to freeze. To avoid this, we placed a projector transparency sheet (Samsill, Fort Worth, TX, USA) between the plywood and glass panes while flies recovered from anesthetization. We put this modified cassette in a 36 °C incubator for 10 minutes after loading the flies then removed the transparency sheets and placed the cassette in the flipping robot. Flies rarely still exhibited freezing behavior. Flies that were stationary (a displacement of less than 1/3 mm between subsequent frames) for more than three quarters of the length of the experiment were excluded from analysis. This criterion excluded less than 10% of flies run in the screen.

Unless otherwise stated, flies were illuminated from above and below by green LEDs (520–530 nm) during behavioral experiments. IR LEDs (850 nm) were used to backlight flies for our video recordings, allowing us to record fly behavior without green illumination. All behavioral experiments lasted 30 minutes and were recorded using a monochrome camera (DMK37-BUX290, The Imaging Source, Bremen, Germany) and an IR pass filter (part FGL715S, ThorLabs, Newton, NJ, USA). Non-mirror experiments were recorded in 1080p at 30 Hz, while mirror experiments were recorded in 1080p at 120 Hz. The videos were compressed and saved using Open Broadcaster Software (OBS).

### Behavioral analysis

All non-mirror behavioral experiments in this study were analyzed using a custom pipeline in Matlab (Natick, MA). A detailed description of the Matlab analysis pipeline is provided in the GitHub repository.

The pipeline first processed the video to find the frames during which the flipping robot rotated the cassette. Once segmented into individual trials (eight second periods during which the flipping robot is stationary), the median image was computed for each trial and subtracted from all frames of that trial, generating a background-subtracted video. This background-subtracted video was binarized using a dynamic threshold followed by a single pixel erosion to remove pixel noise. A MATLAB blob detection algorithm identified individual flies. Since only the flies moved during a trial, they were the only sizeable blobs remaining after background-subtraction.

To analyze the behavior, the user identified key points of the corridors to determine corridor boundaries. The corridor was then segmented into compartments corresponding to the different gaps in the corridor. Then, the fly within each corridor was assigned a unique fly ID that persisted across each frame of the video. If multiple fly-like blobs were identified within a single corridor for a given frame, those blobs were not labeled for that frame (this occurred in less than 1% of frames). The position of the centroid of each blob was extracted for each frame, and an ellipse was fit to each blob. The code determined the heading direction of the fly and assigned the focus of the ellipse in the heading direction as the head of the fly. Based on the position of the centroid, the pipeline automatically determined which corridor compartment the fly was in at each frame. By tracking the transitions between compartments, the code automatically labeled the gap encounter behaviors exhibited at each gap. A detailed diagram of compartment transitions and how they correspond to gap encounter behaviors is provided in the documentation in the GitHub repository.

Although our dataset contained both upward- and downward-directed gap encounters, there were more upward-directed encounters and we restricted our analyses in this work to those encounters that started with the fly vertically below the gap.

This strategy effectively classified some gap encounter behaviors, but it could not distinguish gap crossings from glass circumventions. We trained a neural network to make this distinction from a ground truth data set labeled by human observers. At all four gap sizes, we hand-labeled 1000 events in which the fly moved from one side of the gap to the other as either “gap crossing” or “glass circumvention”. In ∼13% of the cases, it was difficult for the observer to determine whether a fly performed a gap crossing or a glass circumvention. In these cases, the observer still assigned a label, but also additionally labeled the event as ambiguous. The inputs to the neural network were 15 subsampled frames of 100x75 pixels centered on the gap within the second before and after the gap encounter event. The neural networks were three layer, fully-connected convolutional image classifier networks. A standard five-fold cross-validation scheme was used with 20% of the data held out of the training and used as a test set. We trained one network to classify each event as either a gap crossing or glass circumvention and trained a second network to classify each event as a gap crossing or glass circumvention while simultaneously assigning an ambiguity score. The latter network performed better than the former (**Fig. S1E**), so we used it to classify all the data in the MATLAB analysis pipeline. In order to err conservatively, we chose to classify all events labeled as ambiguous by the neural network as glass circumventions. We swept the threshold for what was considered ambiguous (**Fig. S1E–G**) and chose a value of 0.5. This strategy classified approximately 11% of gap traversals as ambiguous (and thus coded as a glass circumvention), while outperforming the other trained network that ignored ambiguity. The classifier-labeled percentages are well within the hand-labeled distribution (**Fig. S1G**).

DeepLabCut^51^ was used to label fly key points for the mirror experiments (Fig. 3). Four key points were tracked on each fly in the mirror: the very rear, the tip of its head, and both eyes. Two key points were tracked on each fly in the corridor (during mirror experiments): the rear and the tip of the head. Several hundred frames were manually annotated in DeepLabCut to train two tracking models, one for tracking flies in the mirror and one for tracking flies in the corridor.

After training the models, they were used to annotate the rest of the frames in the videos which we then manually corrected in frames with large errors. The gap encounter behaviors for the mirror experiments were all manually labeled, and analyses were only performed on gap crossings. For each frame, we computed the angle between the tracked head and the edge of the opposing gap. We then computed the largest instantaneous change in this angle over the course of each trajectory (Fig. 3H) and the largest angular displacement change over the course of the trajectory (Fig. 3I).

### Screen statistics

We devised a data-driven, conservative statistical scheme to assess the statistical significance of behavioral changes when individual neurons were silenced. First, we followed common practice and compared experimental flies, X-Gal4 > UAS-shi, to each parental control, X-Gal4 > + and empty-Gal4 > UAS-shi. However, these comparisons do not account for potential linear additive effects of the Gal4 and UAS constructs on behavior. We therefore generated an additional, synthetic control to account for such additive effects, similar to synthetic controls used elsewhere^96^, where *R_syn_* = *R_WT_* + Δ*_UAS_* + Δ*_Gal4_*, where *R_WT_* is the mean crossing frequency of wild type flies at each gap size, Δ*_UAS_* is the difference in mean crossing frequency between empty-Gal4 > UAS-shi flies and wild type flies at each gap size, and Δ*_Gal4_* is the difference in mean crossing frequency between X-Gal4 > + flies and wild type flies at each gap size. For each silencing experiment, we compared the experimental curve to three controls: the two parental controls and the synthetic control. To make the comparison, we computed p-values using a two-sided Wilcoxon rank sum test for comparisons to parental controls and a two-sided *z*-test for comparisons to the synthetic control for each of the four gap sizes and for the mean difference across all gap sizes. For each control, we assigned the smallest of these five p-values. We considered the control with the largest p-value to be the closest control, and the p-value from the closest control became the uncorrected p-value for that silencing experiment. We then Holm-Bonferroni corrected these across all the Gal4 drivers to obtain the final p-value for each experiment. That correction was for 200 comparisons, equal to the 40 Gal4 drivers examined times the five comparisons made in each of the closest controls. We only considered a Gal4 driver to be a hit if p < 0.05 after Holm-Bonferroni correction, which indicated that the experimental group was statistically significantly different from all three controls.

Driver lines in the screen targeting specific neuron types were all published elsewhere except for the line putatively targeting LPLC4. We are unaware of any published drivers for that cell type, but when viewed in Neuronbridge^93^, that driver is a close match to LPLC4, with LC22 also showing some degree of match. Because we did not observe a phenotype with this Gal4, we did not follow up on validating this driver line.

### Statistical tests for behavior

Each fly was considered an independent sample for statistical purposes in this study, and all means and standard errors were computed over flies. A two-sided Wilcoxon rank sum test was used to test for statistically significant differences between experimental genotypes and parental controls in all behavioral experiments. A *z*-test was used to test for statistically significant differences between experimental genotypes and synthetic controls in all behavioral experiments. All statistical tests in the behavioral screen (Fig. 4**, S4**) were multiple comparison corrected using the Holm-Bonferroni method. In the mirror experiments, we treated each crossing event as a sample when assessing the maximum angular velocity and maximum angular displacement of the far side of the gap (Fig. 3). We used a two-sided Wilcoxon rank sum test to test for statistically significant differences between the three parallax sources in the mirror experiments.

### Derivation of ϕ̇ equation for parallax simulations

Consider a point (*x*′, *y*′) in a stationary reference frame *O*′ and a second reference frame *O* that is translating with velocity *v_x_x̂′* + *v_y_ŷ′* and rotating with velocity −*ωẑ′*. The instantaneous relationship between the coordinates of the two reference frames is:

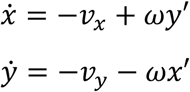

It is also useful to parametrize everything in terms of polar coordinates since we are considering rotations. In polar coordinates, the point (*x*′, *y*′) is written as (*D*′, *ϕ*′), where:

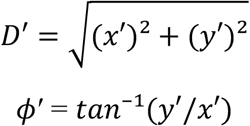

The quantity of interest is *ϕ̇*, which corresponds to the angular velocity of a feature in the visual field of a fly walking along an arc. Orienting the fly in the +x-direction and assuming no crab-walking (*v_y_* = 0), *ϕ̇* is computed as follows:

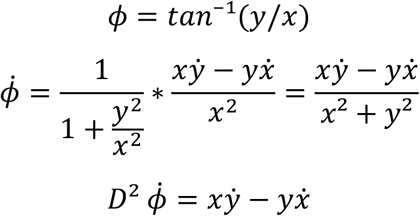

Using the relationships *ẋ* = −*v_x_* + *ωy* and *ẏ* = −*v_y_* − *ωx*, and substituting *v_y_* = 0:

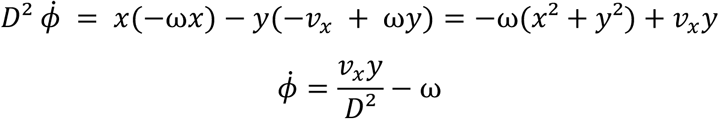

Using the relationship between angular and linear velocity along an arc (*v* = ω*R_c_*)

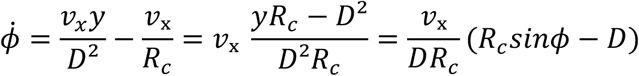

To find points with no motion in the visual field (the dashed purple circle in Fig. 6B), set *ϕ̇* = 0:

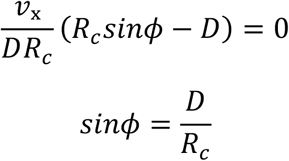

which is the equation of a circle of radius 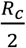 centered at 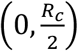 in polar form.

### Geometrical simulation

To better understand the relationships between angular velocity across the eye and relative distance, we performed a geometrical simulation using the equations above (**Fig S6**). We first simulated a fly’s instantaneous trajectory by drawing from a distribution of walking speeds and turning speeds, and simulated an environment by sampling pairs of objects from a distribution of object distances, *D*_1_ and *D*_2_ (**Fig. S6A**), with *D*_2_ always signifying the more distant of the two objects. Given the distributions, we sampled a wide region of space parameterized by the distances normalized by the radius of curvature, *R_c_* (**Fig. S6B**). In this space, the different quadrants correspond to different combinations of front-to-back and back-to-front motion for each object in the pair. For each object, we computed the angular velocity *ϕ̇*, so that we could then plot the joint distributions of *D*_1_/*D*_2_ and *ϕ̇*_2_/*ϕ̇*_1_ for objects in which both were moving front-to-back, both back-to-front, or object 1 moving front-to-back and object 2 moving back-to-front (**Fig. S6C-E**). When next wanted to know how much we could predict about *D*_1_/*D*_2_ from *ϕ̇*_2_/*ϕ̇*_1_. To do this, we created a non-parametric model to predict *D*_1_/*D*_2_ from *ϕ̇*_2_/*ϕ̇*_1_. We first placed the ratio of the angular velocities into 100 equally spaced bins in log-space (as in **Fig. S6C-E**). For each bin, we computed the mean of *D*_1_/*D*_2_, equal to the expected value of *D*_1_/*D*_2_ conditioned on *ϕ̇*_2_/*ϕ̇*_1_. Using these means, we calculated what fraction of the variance in Log_10_ *D*_1_/*D*_2_ was accounted for by this model, generating a coefficient of determination for each of the three sets of motion directions. That coefficient of determination is plotted in Fig. 6C. It depends quantitatively on model parameters and on whether the analysis is done in log or linear space, but qualitatively it consistently shows that when both near and far objects are moving front-to-back, the angular velocity ratio allows better prediction of the distance ratio than in the other cases.

Each parameter for the simulation was drawn independently. The rotational velocity (*ω*) for the fly was drawn from a Gaussian with mean 0°/s and standard deviation 150°/s. The walking speed (*v*) was drawn from a Gaussian with mean 20 mm/s and standard deviation 5 mm/s. Walking speed and rotational velocity distributions are not very different from measured distributions ^72,73^. We only considered flies with walking speeds above 5 mm/s. The objects were drawn from uniform distributions between 0 and 25 mm for *D*_1_ and between *D*_1_ and 50 mm for *D*_2_. We performed 10^8^ of these joint drawings to perform the calculations in **Fig. S6** and Fig. 6C.

### Two-photon imaging

Imaging experiments were performed as previously described^63^ on cold anesthetized male flies expressing GCaMP6f in LC15. Flies were head-fixed onto a stainless-steel shim with UV-curable epoxy. The mouth parts were fixed to minimize brain movement, and the forelegs were removed to prevent interference with visual stimulation. The right optic lobe was surgically exposed by removing the posterior cuticle and fat tissue and was submerged in oxygenated sugar-saline solution^63^. The fly was left for 20 minutes before activity was imaged during visual stimulus presentations using a two-photon microscope (Scientifica, Clarksburg, NJ, USA) equipped with a 20x water immersion objective (Olympus, Waltham, MA, USA). Excitation was provided by a femtosecond Ti-sapphire laser (Spectra-Physics, San Jose, CA, USA) at 930 nm with the power set to < 40 mW. Fluorophore emissions into the photomultiplier tube were filtered by a pair of 512/25 filters (Semrock, Rochester, NY, USA). Fluorescent signals were recorded at 8.46 Hz with ScanImage software^97^.

### Stimulus generation

Visual stimuli were presented at a frame rate of 180 Hz on a virtual cylinder^63^ surrounding the fly using a DLP projector (Texas Instruments, Dallas, Texas, USA). The stimuli were projected onto panoramic screens subtending 270° azimuth and 69° elevation of the fly’s visual field, with a mean screen intensity of ∼70 cd/m^2^. To minimize stimulus bleed-through into the PMT, output from the projector was passed through 565/24 and 560/25 filters (Semrock) in series. The virtual cylinder was pitched forward 45° to approximately account for the pitch of the fly’s fixed head.

All stimuli contained a 50%-contrast random binary checkerboard background with 8° square checks. Each stimulus presentation was preceded by an interleave epoch consisting of 5 seconds of stationary random checkerboard background, with the background pattern retained from the end of the previous stimulus presentation. For stimuli with a moving bar (**Fig. 5E, 5F, 5I, 5J, 6D–G, S6F–K**), a black (–100%) contrast, 8°-wide moving vertical bar was presented. For instances in which both a moving bar and moving background were presented (**Fig. 5I, 5J, 6D– G, S6F–K**), the background began moving for 1 second before the bar initiated its trajectory around they fly, starting from directly behind the fly. In the case of stimuli with a moving vertical bar (**Fig. 5E, 5F, 5I, 5J, 6D–G, S6F–K**), the duration was such that the bar underwent exactly one 360° rotation. In the case of stimuli with only a moving background, the duration was always one second.

### Imaging data analysis

ROIs were defined manually within LC15 dendrites and ΔF/F for each ROI was computed as previously described^63^. Baseline fluorescence (F) for each ROI was obtained by fitting a decaying exponential *Ae^-τ^* to the time-averaged fluorescence for each interleave epoch, with an individual *A* fit to each ROI and the same *τ* fit to all ROIs in a single recording. To obtain ΔF, the exponential fit was subtracted from the ROI-wise time trace values.

Each stimulus was presented four times per fly. Within each ROI, the responses to repetitions of the same stimulus were averaged. Then, the responses across ROIs for the same stimulus were averaged to obtain an individual fly-wise mean response trace. To obtain group statistics, these individual fly-wise values were used in calculating group means and standard errors of the mean (SEMs). Mean response to a stimulus was calculated by averaging ΔF/F over the full-width half-max of the fly’s peak response (**Fig. 5F, 5H, 5J, 6F, 6G, S6G, S6H, S6J, S6K**). In the case of a visual stimulus with only a background (**Fig. 5G, 5H**), mean response was calculated averaging over one second of stimulus presentation.

### Statistical tests for imaging

All pairwise comparisons were conducted via Student’s *t*-test with a Holm-Bonferroni correction (**Fig. 5F, 5H, 5J**). We evaluated the decoding predictivity of neural response for each of three target decoded features (bar velocity, background velocity, or the ratio of the two). We trained three different models on the same data (**Fig. 6I–K**) to predict stimulus conditions for the three target features, and calculated the coefficient of determination (R^2^) for each decoder. We binned the normalized responses of each fly into *k =* 10 bins (each 0.1 wide). To calculate R^2^, the ratio of the residual sum of squares to total sum of squares was used:

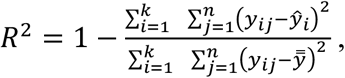

where *i* corresponds to a normalized neural response bin and *j* corresponds to an individual fly. The dependent variable, *y,* was the stimulus condition of a given target feature to be predicted by the response bin *i,* such that *y_ij_* is the stimulus condition corresponding to the LC15 response bin *i* in fly *j*. Here, *ŷ_i_* represents the average stimulus condition for a given response bin *i*, and *y̿* represents the grand average of the stimulus condition *y*.

To avoid assumptions of normality in our estimates of R^2^, we calculated bootstrapped distributions with 10,000 iterations to obtain 95% confidence intervals (**Fig. 6I–L**). For pairwise comparisons between the decoding predictivity of bar velocity, background velocity, and the ratio of the two velocities in paired data (**Fig. 6I–K**), where all comparisons were made within the same set of flies, *p*-values were computed as the two-tailed percentile of bootstrapped values for which the difference in model predictivity (|Δ*R*^2^|) was greater than the measured difference^98^. When making pairwise comparisons across different samples of flies (**Fig. 6L**), a bootstrapped null distribution of the difference in R^2^ was computed with resamples drawn from two groups pooled together, and *p*-values were calculated from the two-tailed percentile in the null distribution greater than the empirical difference in R^2^ ^98^.

